# Persistence as an optimal hedging strategy

**DOI:** 10.1101/2019.12.19.883645

**Authors:** Alexander P Browning, Jesse A Sharp, Tarunendu Mapder, Christopher M Baker, Kevin Burrage, Matthew J Simpson

## Abstract

Bacteria invest in a slow-growing subpopulation, called persisters, to ensure survival in the face of uncertainty. This hedging strategy is remarkably similar to financial hedging, where diversifying an investment portfolio protects against economic uncertainty. We provide a new theoretical foundation for understanding cellular hedging by unifying the study of biological population dynamics and the mathematics of financial risk management through optimal control theory. Motivated by the widely accepted role of volatility in the emergence of persistence, we consider several novel models of environmental volatility described by continuous-time stochastic processes. This allows us to study an emergent cellular hedging strategy that maximizes the expected per-capita growth rate of the population. Analytical and simulation results probe the optimal persister strategy, revealing results that are consistent with experimental observations and suggest at new opportunities for experimental investigation and design. Overall, we provide a new way of conceptualising and modelling cellular decision-making in volatile environments by explicitly unifying theory from mathematical biology and finance.

## Introduction

Shortly after the clinical introduction of penicillin, Bigger noticed a resistance in a small subpopulation of *Staphylococcal pyogenes* (1). The resistant cells, termed *persisters*, are a genetically identical, slow-growing, phenotypic variant. Bacteria invest in persisters to ensure survival: persisters are less proliferative than regular cells in a nutrient rich environment but can withstand adversity (2). These strategies are akin to financial hedging, where diversifying an investment portfolio protects against economic uncertainty (3–6). This similarity is widely acknowledged: parts of the biology and ecology literature refer to this phenomena as *bet-hedging* (2, 5, 7–9). In this work, we introduce the concept of *cellular hedging*, where we explicitly model bacterial persistence using techniques from mathematical finance, including stochastic differential equations (SDEs) and stochastic optimal control theory.

Bacterial persistence poses a significant clinical challenge and is highly advantageous to bacteria. For example, persisters are less sensitive to antibiotic treatment (10–13), are undetectable in routine clinical tests (14), and are thought to be responsible for the formation (15) and incurability (16) of many infections. Antimicrobial treatments will, therefore, benefit from an understanding of how persisters arise and function (17). The similarity between bacterial persistence and financial hedging strategies suggests a new pathway to investigate persister dynamics. In this study, we provide a novel, quantitive understanding of persister strategies using techniques from financial mathematics and stochastic optimal control theory. Persister production is known to depend on the environment (10, 18, 19) and our mathematical modelling approach allows us to unearth how various optimal persister strategies depend upon environmental volatility.

Persisters can be revealed experimentally by disinfecting a population of *Escherichia coli* (*E. coli*) with ampicillin, which targets proliferative cells (10). The initially rapid reduction in population size eventually slows, revealing a small subpopulation that is less sensitive to the treatment. Experimental investigations are complicated by the extreme scarcity of persister cells: typically less than 1 in 10^5^ cells are persisters in wild type *E. coli* (20). For subpopulations of this size, stochastic effects are significant (21, 22).

In producing persisters, bacteria allocate resources to hedge against environmental volatility for the purpose of survival. We assume that these processes occur in much the same way that a financial investor hedges a portfolio to protect from, and take advantage of, economic volatility (23). The archetypal example of portfolio diversification in finance is Merton’s portfolio problem (MPP) (3). Here, an investor allocates a fraction of their wealth in a high-yield volatile asset, such as stocks; and a low-yield stable asset, such as government bonds. Analogously, we suppose that bacteria have evolved mechanisms to regulate an allocation of their total population as proliferative although susceptible, and some as persisters. In the finance problem, market volatility is modelled such that the underlying price of each asset can be described as an Itô SDE driven by Wiener noise (3, 24), similar to models of noisy exponential growth proposed for biological problems (25). More complicated models of market volatility have been extensively explored in the finance literature, such as those that incorporate Poisson jump noise to describe market shocks (26). In Merton’s original work, stochastic optimal control theory (24) is used to maximize the investors wealth by modelling the proportion allocated to each asset as a *control*. Merton revealed that it is not advantageous to possess the low-yield asset in the absence of uncertainty, or when the growth of the high-yield asset is deterministic. MPP revolutionized the field of mathematical finance, and many of the ideas in MPP formed the basis of Merton’s later work on options pricing with Black and Scholes (27, 28) that led to the the 1997 Nobel Prize in Economics.

Current mathematical models of persistence typically describe environmental volatility with a growth rate that transitions through finitely many states (such as a growth and stress state). Transitions between these states are assumed to occur either periodically (8, 21, 29) or stochastically (4, 5, 30). Many existing studies maximize some measure of the long-term growth rate of the population and probe the environmental conditions under which persistence is advantageous using, for example, Lyapunov exponents (30). A limitation of these existing modelling approaches is their inability to capture more general models of environmental volatility. We leverage an explicit connection with mathematical finance to study optimal cellular hedging strategies using stochastic optimal control theory (24, 31). This approach has several advantages. First, we can probe cellular hedging strategies under more complex models of environmental volatility. Second, we can model the emergence of an environment-dependent persister strategy.

In our study, we first model persistence in a population where the growth rate is subject to continuous stochastic fluctuations in the form of Wiener noise (22, 24, 32). We posit that, in many cases, this is a more appropriate model of environmental volatility than the discrete transitions, or shocks, that are currently commonplace in mathematical models of persistence (8, 21, 29, 30). Through this model, we draw a direct connection with mathematical finance and demonstrate how MPP can be applied directly to the biological problem. Next, we expand on this simplistic model through an environment-dependent hedging strategy under more complex models of environmental volatility. We include in our analysis a model that uses Poisson jump noise (31), to capture the aforementioned existing models of environmental volatility (4, 5, 8, 21, 30) where transitions in the growth rate occur due to, for example, shocks. Our modelling framework is readily extensible to any form of environmental volatility that can be described using an Itô SDE.

Our goal is to develop a new framework for studying cellular hedging in response to environmental volatility by unifying the study of biological population dynamics with techniques financial mathematics. It is for this reason we use the term *cellular hedging*, instead of the term *bet-hedging* (2, 5, 7–9). Despite the established importance of volatility in elucidating bacterial persistence, we find there is currently a scarcity of methods available to describe cellular hedging strategies under environmental volatility. Our mathematical framework demonstrates the importance of considering more complex models of environmental volatility, and we lay theory to complement future experimental studies that probe emergent persister strategies in response to continuously varying stochastic environments. Our new model and approach leads to mathematical results that are consistent with observations from several existing experimental studies, and provides new insights into bacteria dynamics in the face of environmental volatility.

## Methods

### Stochastic model of bacteria growth

We describe population-level bacteria dynamics in a volatile environment with a system of SDEs driven by Wiener noise (22, 25, 32). This choice of model can be thought of as a bridge between discrete Markov models (21, 23), and deter ministic ordinary differential equation (ODE) models (10). In addition, our population-level model of bacterial persistence complements stochastic gene expression models that describe the regulation of persister strategies (6, 33). Further, our modelling approach can be generalized to can capture many other forms of environmental noise. For example, we model the dynamics of bacteria in a volatile environment by coupling the growth rate to a stochastic process representing the environment (34).

We model exponential growth in a population composed of regular (non-persister) cells, *r_t_*, and persisters, *p_t_*. Here, a subscript *t* indicates that each stochastic processes depends on time. The net growth rate of regular cells is a stochastic process with expectation *μ_t_* and amplitude *σ*. The quiescence of persisters manifests as slow metabolic activity, so we assume the expected growth rate of persisters is some small proportion, *ɛ* ≪ 1, of regular cells. The net persister growth rate is, therefore, a stochastic process with expectation *ɛμ_t_* and amplitude *η* = *εσ*. In the absence of subpopulation switching, these dynamics give rise to the system of differential equations (25, 32)

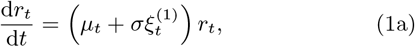

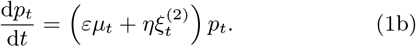

Here, 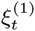 and 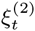 are independent Gaussian white noise processes. The expected growth rate of regular cells is given by *μ_t_*, which may be constant, or itself a stochastic process. We interpret Equation 1 in the Itô sense for two reasons. First, the properties of the Itô integral make it well suited to models in population biology (25, 35). Second, Itô SDEs are widely applied in mathematical finance, including in MPP (3), where the property 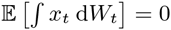, in the Itô sense, is vital.

The current literature classifies two types persister production in a population: variable and environment dependent, commonly referred to as Type I; or constant, commonly referred to as Type II (10) (Fig. 1). It is understood that constant production of persisters, which results in an approximately constant proportion in a growing population, may be regulated by stochastic gene fluctuation on a single cell level (36–38). In addition, a growth feedback mechanism (39) — possibly regulated by quorum sensing (40) and intercellular signalling (41) — may enable cells to respond and vary the persister production rate. In our model, regular cells switch to persisters at a rate of *u* + *ϕ_t_*, where: *u* ≥ 0 is the constant rate (10); and *ϕ_t_* ≥ 0 is the variable, environment-dependent, rate. Persisters revert to regular cells at a constant rate *v* ≥ 0. Our model can, therefore, describe both Type I and Type II strategies.

**Fig. 1.**
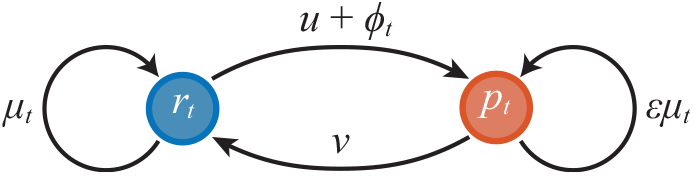
Schematic of the bacteria growth model with regular cells, *r_t_*, and persisters, *p_t_*. Switching from persisters to regular cells is constant, *v*. Switching from regular cells to persisters is taken to be the sum of a constant rate, *u*, and a variable rate, *ϕ_t_*, that depends upon environmental volatility.

Incorporating subpopulation switching into the growth equations (Eqs. 1) yields

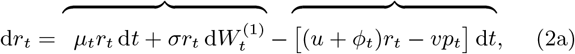

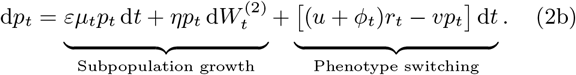

Such a system of equations is said to be in the form of multiplicative Wiener noise (31). We note that in removing variability from the model by setting *σ* = *η* = 0, we recover the ODE model of Balaban *et al.* (10). Many other choices for a stochastic population dynamics model exist (22), such as models that consider intrinsic noise caused by subpopulation switching (42).However, the focus of the current work is on fluctuations in the environment and growth rate, which we assume to be independent of the population size.

We find it natural to consider the variable transformation

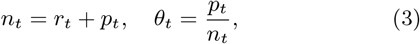

such that *n_t_* represents the population size and *θ_t_* represents the proportion of persisters. Following Itô’s lemma (24), the transformed state equations are

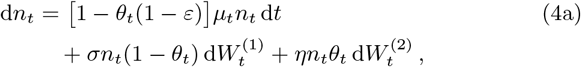

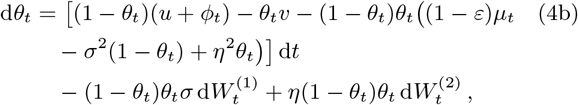

revealing that the dynamics of the persister proportion, *θ_t_*, are independent of the population size, *n_t_*.

### Stochastic model environmental volatility

We model environmental volatility by assuming *μ_t_* = *m*(*ζ_t_*), where *ζ_t_* is a stochastic process that represents a volatile environment. There are many appropriate choices for *m*(·) and *ζ_t_*, but we focus our analysis on three environments, samples paths of each shown in Fig. 2*a–c*. These are

**Fig. 2.**
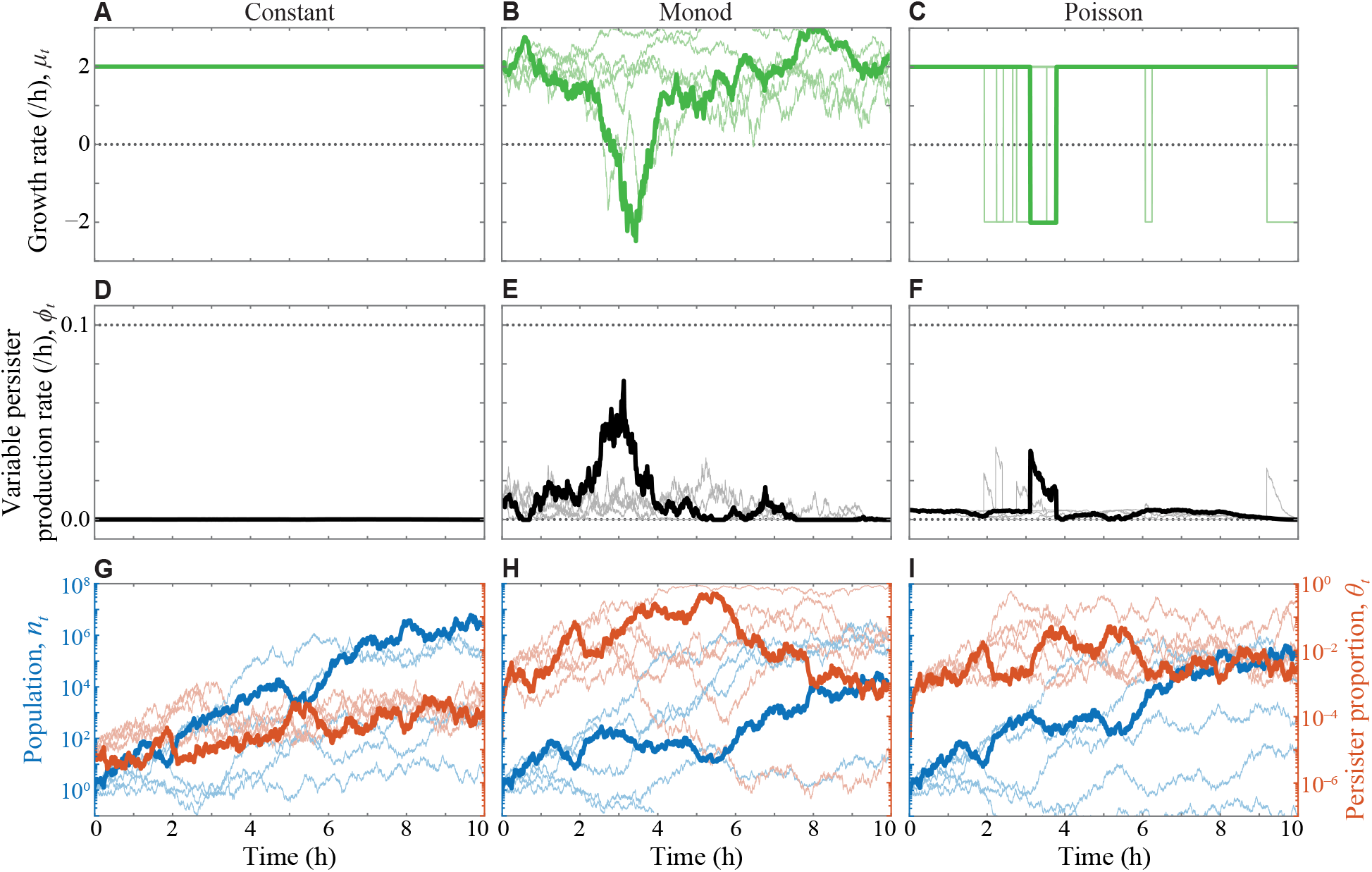
Optimal persister production in cell populations under various types of environment. (a)-(c) Growth rates, *μ_t_*, sampled from each environment to simulate bacteria growth in the rest of each column. (d)-(f) The optimal variable persister production rate, 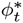.(g)-(i) The population, *n_t_*, in multiples of the original population(blue, left scale) and persister proportion, *θ_t_*, (red, right scale), for a single realisation of the model. The seeds used to generate the Wiener processes 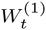 and 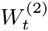 are identical for all environments. Also shown are five additional, independent, realisations (semi-transparent).

#### 1. Constant

(Fig. 2*a*) We set *m*(*ζ*) ∈ {*μ_G_, μ_S_*}, such that *μ_t_* is constant. Here *μ_G_* represents a colony during growth; and *μ_S_* represents a colony under stress or antimicrobial treatment. For numerical results, we choose *μ_G_* = 2 h^−1^ and *μ_S_* = −2 h^−1^ to match experimental data for *E. coli* during growth and ampicillin treatment (10).

#### 2. Monod

(Fig. 2*b*) Monod kinetics are commonly used to model the growth of bacteria (43–45), and feature dynamics with an asymptotic upper bound on the growth rate, but no lower bound. We describe this environment by a mean-reverting Ornstein-Uhlenbeck process (24) and couple the growth rate using a Monod equation (43),

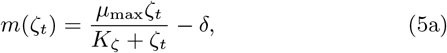

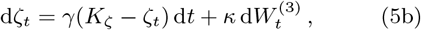

where 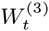 is a Wiener process independent of both 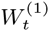 and 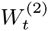. Here, *ζ_t_* experiences fluctuations proportional to *κ* and a reversion force to the state *ζ_t_* = *K_ζ_* of strength proportional to *γ*. The Monod coupling features an asymptotic upper-bound on the growth rate of *μ_t_* = *μ*_max_ and no lower bound. Unfavourable environmental changes have, therefore, a larger effect on the growth rate than those that are favorable, and the growth rate experiences reversion to a growth rate *μ_t_* = *μ*_max_/2 − *δ*, where *δ* represents the natural death rate. For numerical results, we choose *γ* = 0.1 h^−1^, *K_ζ_* = 1, *ζ*_0_ = 1, *κ* = 0.3, *μ*_max_ = 8 h^−1^ and *δ* = 2 h^−1^. For this choice of parameters the growth rate experiences reversion to a growth rate of *μ_G_* = 2 h^−1^, which corresponds to the constant environment. To deal with the discontinuity at *ζ* = −*K_ζ_*, we truncate Eq. 5a so that *m*(*ζ*) = *m*(−0.5) for *ζ* < −0.5.

#### 3. Poisson

(Fig. 2*c*) An existing class of mathematical models describe environmental uncertainty using alternating periods of growth and stress (4, 5, 8, 21, 30). We reproduce this type of environment in a stochastic model by assuming that the environment switches between growth and stress according to a Poisson process. We consider that *m*(*ζ*) = *μ_G_* for *ζ* ≥ 0; *m*(*ζ*) = *μ_S_* for *ζ* < 0; and model the environment as the jump or telegraph process (31)

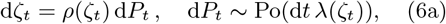

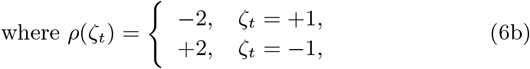

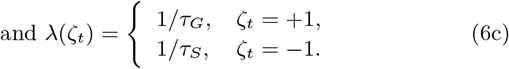

Here, *p_t_* is a Poisson process with intensity *λ*(*ζ_t_*). This formulation allows for a mean time of *τ_G_* in the growth phase, where *μ_t_* = *μ_G_*, and a mean time of *τ_S_* in the stress phase, where *μ_t_* = *μ_S_*. For numerical results in this study, we choose *τ_G_* = 9.5 h, *τ_S_* = 0.5 h, *ζ*_0_ = 1, *μ_G_* = 2 h^−1^ and *μ_S_* = −2 h^−1^. This model of environmental volatility can be readily extended to capture more general forms of Poisson noise leading to discontinuous fluctuations in the environment due to, for example, shocks.

These choices represent environments in which changes happen gradually (Monod environment) or abruptly (Poisson environment), as demonstrated in Fig. 2*b* and Fig. 2c, respectively. For comparative purposes, we demonstrate results using realisations of each environment that are similar, and choose the initial condition for all environments to correspond to the initial growth rate *μ*_0_ = *μ_G_* = 2 h^−1^. In the supporting material, we explore results for two other models of environmental volatility: (4) an Ornstein-Uhlenbeck process without a Monod coupling (Fig. S1d) and, (5) a Duffing oscillator (Fig. S1e).

### Optimal persister production strategies

Our ideas of *cellular hedging* suggest that a cell population has developed a persister production strategy 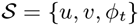 that is optimal in some way. Mathematically, we define optimality as the strategy that maximizes some *fitness measure*, which we now construct. We assume that there is no explicit cost to producing persisters with constant rates *u* and *v*; but that there is a quadratic running cost to produce persisters with a variable rate *ϕ_t_*. The cost of applying a non-zero *ϕ_t_* accounts for the sensing mechanisms that cells must use to respond to the environment.

We choose a fitness measure, commonly referred to as a *payoff* in optimal control theory, as

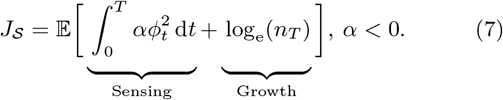

Here *T* denotes a terminal time, so that the maximisation is carried out on the interval *t* ∈ [0*, T*]; *α* < 0 characterises a trade-off between growth and operating the sensing mechanisms required to vary the persister production rate; and the expectation is taken with respect to the stochastic processes governing subpopulation growth, *n_t_*, and the environment, *ζ_t_*.In this work, we consider a finite terminal time since we model an exponentially growing population. It is not obvious how to incorporate stationary phase dynamics into the stochastic environment model. Future work is needed to explore the infinite time-horizon problem in conjunction with environmental volatility and a logistic growth term (25). We interpret the terminal time, *T*, as either the duration of the growth phase of the population (10), or the duration of an experiment.Maximising the logarithmic term in Eq. 7 can be interpreted as cells maximising their per-capita growth rate over the interval *t* ∈ [0*, T*], since

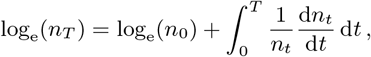

and log_e_(*n*_0_) is constant (46). Our choice of fitness measure corresponds to evolutionary mechanisms that perpetuate highly productive populations (21), meaning cells that implement the optimal strategy carry an evolutionary advantage over those that do not.There are many other choices of fitness, such as resource allocation based on a maximum entropy principle (47). We expect our methodology to carry across to these more complicated choices of fitness measure.

Together with the state equations (Eq. 4), maximising Eq. 7 corresponds to an optimal control problem. We apply Hamilton-Jacobi-Bellman (HJB) optimal control theory (31, 48, 49) to reformulate the optimal control problem as a partial differential equation (PDE) problem. We now review essential elements of HJB optimal control theory.

### Review of optimal control theory

A stochastic control problem may be stated as

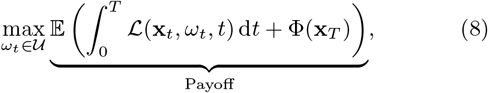

where *ω_t_* is a time-dependent *control* from the set of allowable controls, 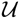. The payoff contains terms 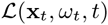, which represents cumulative value (in our case, the sensing mechanisms), and Φ(**x**_*T*_), which represents the terminal value (in our case, the final population size). For a stochastic problem, the payoff involves an expectation, which is taken with respect to the stochastic process **x**_*t*_, representing the sys-tem state. The goal is to find the so-called *optimal control*, denoted 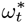, that maximizes the payoff.

The state equations are given by the three-dimensional stochastic process, **x**_*t*_ = (*n_t_, θ_t_, ζ_t_*), governed by the three-dimensional Itô SDE

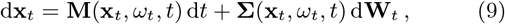

where **M** ∈ℝ^3^ is a vector valued function containing the drift for each state; d**W**_*t*_ ∈ ℝ^3^ is a Wiener process of the same dimension as **x**_*t*_ with independent constituents; and, **ΣΣ**^tr^ ℝ^3×3^ describes the covariance of the Wiener process (superscript ^tr^ denotes the matrix transpose). In our study, **x**_*t*_ comprises the stochastic processes governing the growth of the bacteria population, (*n_t_, θ_t_*), in addition to the independent stochastic process for the environment, *ζ_t_*. We also study a form of **x**_*t*_ that includes Poisson jump noise, and we include details of optimal control theory in this case as supporting material.

To solve the optimal control problem we define a *value function*, *V* (**x***, s*), as the optimal payoff obtainable were the system to be at state **x** at time *s*. That is,

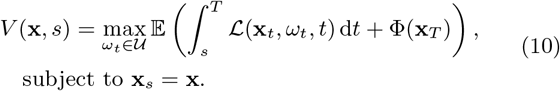

HJB optimal control theory then describes the time-evolution of *V* as the solution of a partial differential equation (PDE) (48, 49), given by

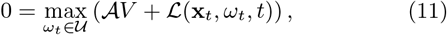

where 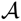 is the stochastic generator for the process governing **x**_*t*_. For the three-dimensional system given by Eq. 9, the stochastic generator is given by

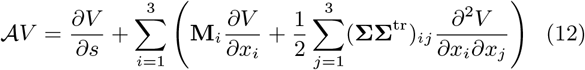

where **M**_*i*_ denotes the *i*th element of **M**(**x**_*t*_, *ω_t_, t*), and (**ΣΣ**^tr^)*ij* denotes the element in the *i*th row and *j*th column of **ΣΣ**^tr^, for **Σ**= **Σ**(**x**_t_, *ω_*t*_, _t_*). Therefore, mixed partial derivatives are only included in Eq. 12 in the case where the stochastic processes are correlated. The idea behind HJB optimal control theory is that *V* (**x***, s*) is known at time *s* = *T*. Therefore, Eq. 11 is coupled to the terminal condition *V* (**x***, T*) = Φ(**x**_*t*_), and solved backwards in time (50).

In certain cases, the argmax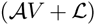 term in Eq. 11 may be found analytically through differentiation. This yields both an expression for the optimal control, 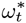; and a three-dimensional, non-linear, PDE coupled to a terminal condition. In the case of MPP, where the volatility appears only as geometric Brownian motion in the state equations for (*nt, θt*), and not in an independent equation for the environment,Eq. 11 has an analytical solution (3). In most cases, however, we are required to solve Eq. 11 numerically. To obtain an optimal trajectory starting at state **x**_0_, we solve the state equations (Eq. 9), coupled to the solution of the HJB PDE (Eq. 11), forward in time using the Euler-Maruyama algorithm (51). Full details of the numerical techniques are provided in the supporting material.

### Numerical methods to solve the SDEs

We integrate the SDE models using the Euler-Maruyama algorithm (51) with time step *h* = 2 × 10^−3^. To compare results between environments, we fix the seeds used to generate the Wiener processes 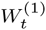 and 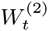 when simulating different models. Full details of the numerical method used to solve the SDEs are given in the supporting material.

### Constant persister production

We first examine a population that can only produce persisters at a constant rate, so we fix the variable rate, *ϕ_t_* = 0. In this case, we only assume the constant environment so *μ_t_* = *μ_G_*.

As *u* and *v* only appear in the optimal control problem linearly, the solution is not finite unless a bound is enforced on *u* and *v* (52). The unbounded problem corresponds to a cell population that is able to move itself instantaneously to anywhere in the state space. To address this, we assume that the constant switching rates *u* and *v* are chosen by the population to control the steady-state persister proportion, which we denote 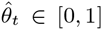. Taking the drift term in the state equation for *θ_t_* (Eq. 4b) to be zero, we see that *u* and *v* are related to 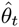 by

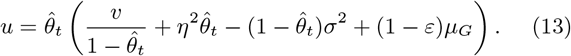

At this point, we note that Eq. 13 is only consistent if 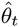 is constant, and we address this shortly.

Allowing 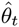 to be a control, the state equation for *n_t_* (Eq. 4a) becomes

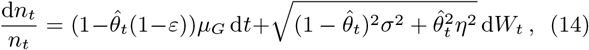

since 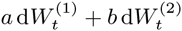 can be considered as 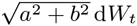 Assuming that 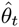 is chosen to maximize the fitness measure (Eq. 7) reveals a problem closely related to MPP (3). The problems differ intrinsically in that an investor is able to reallocate assets within a financial portfolio instantaneously (23) and without cost — though variations of MPP address this— whereas the cell colony comprises finitely many cells that act heterogeneously using finite switching rates to produce persisters.

We apply stochastic optimal control theory (24, 31) and exploit the similarity between this formulation of the persister problem and MPP (3) to obtain an analytical solution, in which the optimal control, denoted 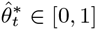, is given by

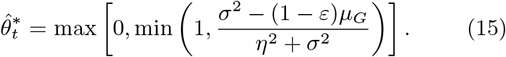

Full details of this analytical solution are given in the supporting material. As is the case with MPP, 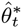 is constant, so the strategy is independent of the current time, the terminal time, and the system state.Solving Eqs. 13 and 15 simultaneously gives 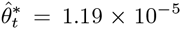 with *σ* = 1.414222 which matches experimental observations for wild type *E. coli* bacteria where *μ_G_* = 2 h^−1^, *ε* = 0, *η* = *εσ* = 0, *u* = 1.2 × 10^−6^ h^−1^ and *v* = 0.1 h^−1^ (10). Here, we note that *σ*^2^ ≈ *μ_G_*, since 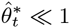.

For numerical results in the rest of this study, we fix *σ*, *ɛ*, *η*, *u* and *v* to the aforementioned values, and set the initial persister proportion *θ*_0_ = 1.19 × 10^−5^. We set *n*_0_ = 1, such that the population is measured relative to the initial population. In practise, these parameters would be obtained by calibrating the stochastic model to experimental data, where we expect *σ*, and, therefore, *u* and *v*, to depend upon the stochastic environment induced by the experimental conditions. Further, setting *σ* = *η* = 0 would recover a model in which *ζ_t_* is the only source of environmental volatility.

### Variable persister production

We now consider a cell colony that is able vary persister production in response to their environment. In this case, *ϕ_t_* ∈ ℝ^+^, can be thought of as a Markovian control (24), or a control that is chosen based on information that includes the current state and time, but does not carry a memory about the past state. For numerical stability, we place an upper-bound on the control such that *ϕ_t_* ≤ 0.1 h^−1^.

To solve the control problem, we define the value function, *V*, by

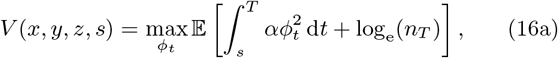

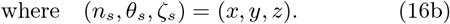

HJB optimal control theory describes *V* with a PDE, expressed in this case as

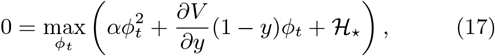

where 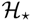 represents a collection of terms independent of the control, *ϕ_t_*. As Eq. 17 is quadratic in *ϕ_t_*, we can carry out the maximisation by setting the derivative to zero to find the optimal control, denoted 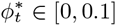, as

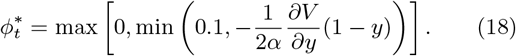

Here, *y* = *θ_t_* is the current persister proportion. Since *y* ∈ [0, 1], an interpretation of Eq. 18 is that the population uses 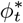 to steer the population toward the optimal persister proportion, where *∂V/∂y* = 0. As environmental triggers only create persisters (we assume the quiescent state of persisters means they are unable to react to the environment), 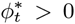, and so 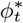 is only active if the current proportion is less than the optimal proportion. Finally, 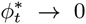 as *α* → −∞, which represents the sensing mechanisms becoming prohibitively expensive.

Substituting Eq. 18 into the HJB equation leads to a non-linear PDE for *V* that must be solved backward in time. Full details of this equation are given in the supporting material. In summary, the variable transformation *x* → log_e_(*x*) removes all terms containing the independent variable *x* (representing the current population size) from the equation. Therefore, we find that the ansatz

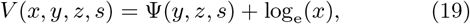

is consistent with the system. This result reveals that the optimal control (Eq. 18) is independent of the population size since *∂V /∂y* = *∂*Ψ*/∂y*. This gives an optimal strategy of the form 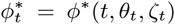. These results suggest that bacteria can implement an optimal variable persister strategy using only information about the proportion of persisters (through, for example, quorum-sensing (40) or intercellular signalling (41)), and the environment (through, for example, a growth feedback mechanism (39)). Therefore, the persister strategy is dependent upon stochastic fluctuations in both the proportion of persisters and the environment.

In Fig. 2 we show dynamics for cell colonies in all three environments for *α* = −100 h. For these results, we solve for Ψ numerically. As expected, we find that unfavorable environments trigger variable persister production (Fig. 2*e,f*), without any additional persister production for the constant environment (Fig. 2*d*). This suggests that a constant persister production strategy is sufficient for an environment with a constant expected growth rate. On the other hand, additional persister production is seen in response to environmental cues for more volatile environments (Fig. 2*e,f*). An interesting result in Fig. 2*e* is that the peak variable production rate under the Monod environment lies before the minimum growth rate. This suggests that the model implicitly incorporates mechanisms that allow cells to respond to features of the stochastic environment, to the extent that they may be able to anticipate environmental changes based upon their own current growth rate.

### Behaviour under unfamiliar environments and antimicrobial treatment

By assuming that cells monitor their growth rate, *μ_t_*, we can explore how bacteria that are specialized to one environment (i.e. implement the optimal persister strategy) behave under an unfamiliar environment. We couple the growth rate to the optimal control by considering 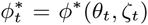 where *ζ_t_* = *m*^−1^(·) denotes the inverse of the growth rate coupling function *m*(*ζ*). In other words, the cells measure the current environment state using the growth rate.

Persisters are revealed experimentally by exposing the population to antibiotics and monitoring the population size (10). We simulate this process to examine how bacteria specialized to each environment behave when exposed to a constant expected decay rate. Simulation results in Fig. 3*a*, for a population that can only produce persisters at a constant rate, are similar to experimental results for wild-type *E. coli* (10). In comparison, we find that colonies that implement a strategy optimal under a more volatile environment, such as the Poisson and Monod environments, produce persisters using the variable rate (Fig. 3*c*) reach saturation of persisters more quickly (Fig. 3*b*) and have a higher long term population (Fig. 3*a*). These qualitative observations suggest that high persistence bacteria strains (10, 21) could arise in response to highly volatile environments. In the supplementary material we apply the same methodology to simulate bacteria specialized to each environment under other, unfamiliar, environments. These results demonstrate the potential of our work to model, for example, how an optimal persister strategy interacts with therapeutic interventions such as drug sequencing (53, 54).

**Fig. 3.**
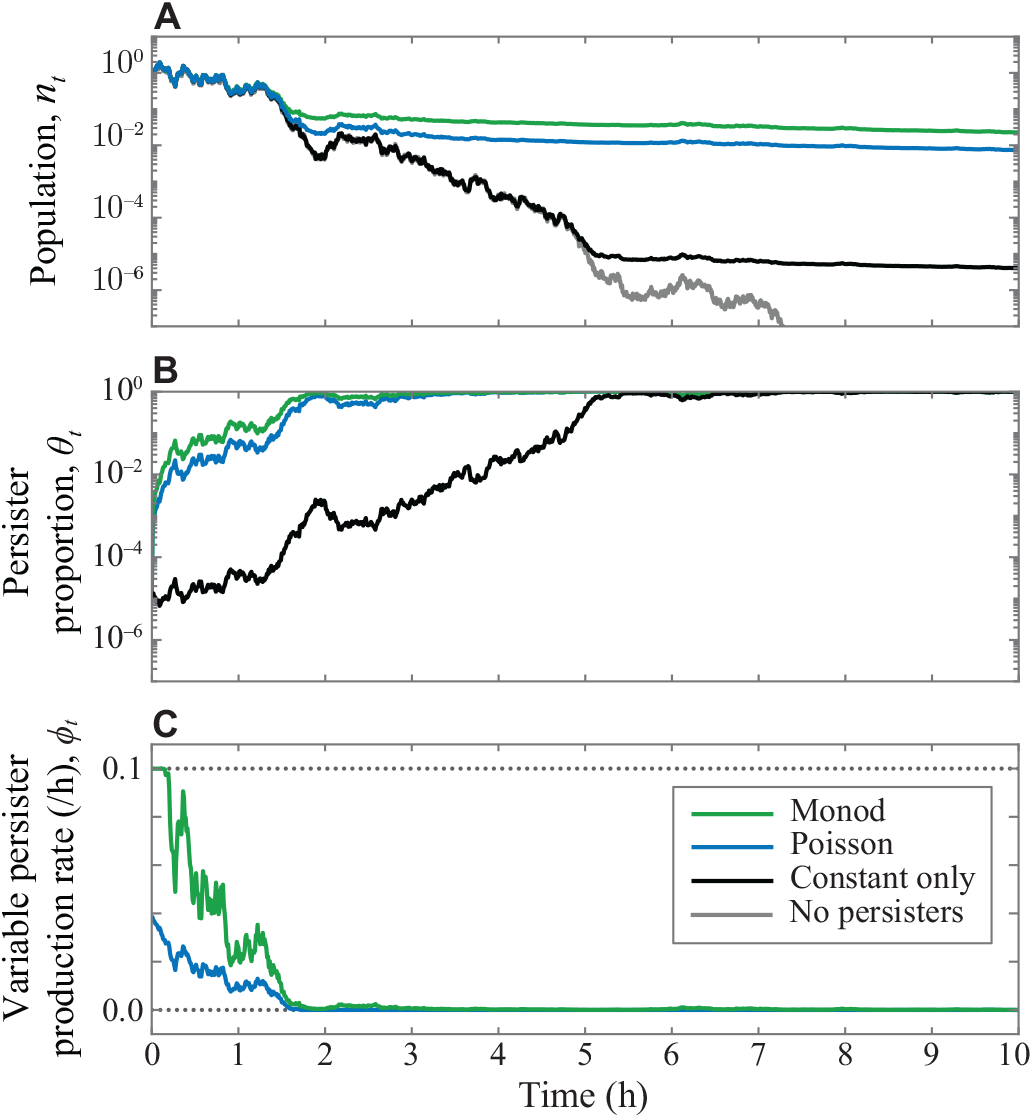
Single realisations of the model showing the behaviour of a cell population under antibiotic treatment. (a) Shows the population, *n_t_*,in multiples of the original population;(b) shows the persister proportion, *θ_t_*; and, (c) shows the variable persister production rate, 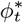. In all cases, the cells experience a growth rate *μ_t_* = 2 h^−1^ and implement: the variable strategy optimal under either the Monod environment (green) or the Poisson environment (red); constant switching only optimal under the constant environment (black); or do not produce persisters (grey).

## Discussion

We describe persister dynamics with a stochastic model and introduce the idea of *cellular hedging* to study an optimal persister production strategy. By applying cross-disciplinary ideas from mathematical finance to the persister problem, we provide several analytical and simulation results that elucidate bacterial persistence under several novel models of environmental volatility.

Our results conform to the consensus view that it is detrimental to produce persisters in the absence of volatility (9, 21, 23, 55).In examining the optimal strategy for constant persister production, we reveal the proportion of persisters that must be maintained to maximize the expected growth of the population (Eq. 15). Removing volatility from the model by setting *σ* = 0 recovers the deterministic model of Balaban *et al.* (10) and reveals an optimal persister proportion of zero. Therefore, our model predicts that producing persisters in the absence of environmental volatility is suboptimal and, therefore, leads to a lower expected per-capita growth rate. In contrast, the straightforward assumption of a growth rate driven by Wiener noise reveals a small proportion of persisters (approximately 1 in 10^5^) — consistent with experimental observations of Balaban *et al.* (10) — that maximizes the expected per-capita growth rate. Our results contrast with previous mathematical studies that assume external factors, such as antibiotic treatment, to explain persistence (21).

Compared with hedging strategies in finance, that often rely on complex, market-level information, a cellular hedging strategy must be limited in complexity. A key result of the direct comparison between bacterial persistence and MPP is to reveal a constant optimal proportion of persisters that should be maintained in an environment with a constant expected growth rate. This result is significant for the persister problem as, unlike in MPP, the cell population cannot directly control the proportion of persisters. This key difference between the biological and financial hedging problems is widely understood in the literature (23). However, we show how a population can maintain a constant *expected* proportion of persisters, regulated by constant switching propensities, *u* and *v*.

The mechanisms that enable cells to implement environment-dependent cellular hedging strategies are unknown (56), but must come at a cost to the cells—in terms of additional genetic machinery—and rely on the limited information available to individual bacteria. Our study reveals an optimal variable persister production strategy that depends only on the time, persister proportion and the environment, but not on the the population size. Additional results in the supporting material (Fig. S6) indicate that these strategies become time-independent far from the terminal time, *T*. If an infinite terminal time problem was studied, necessarily with a growth model that incorporated crowding effects (such as logistic growth), we expect the optimal strategy to be independent of time. Furthermore, the optimal environment-dependent strategy revealed in Eq. 18 is not complex. Analysis of Eq. 18 suggests that bacteria increase persister production when the persister proportion for a given growth rate is less than optimal. This kind of information could be genetically encoded as an evolutionary adaptation to a particular environment. The parameter *α*, therefore, specifies the rate at which the population is able to respond to changes in the environment: if variable switching is expensive (*α* ≫ 1), the population will respond more slowly than if variable switching is relatively cheap.These results are surprising as optimal control theory only provides the strategy that maximizes the payoff, and does not necessarily enforce any level of complexity in the cellular hedging strategy.

In Fig. 3, we simulate a persister producing bacteria colony under antibiotic treatment, demonstrating the distinct advantage persisters afford bacteria (10, 14, 17). Our stochastic model of persister production can be used to improve the efficacy of antimicrobial therapies using optimal control from an optimal treatment perspective (52, 57–59). Furthermore, recent experimental and mathematical work examines the potential of so-called *evolutionarily informed therapy* or *drug sequencing* (53, 54, 60), where a sequence of drugs is administered to sequentially induce susceptibility and overcome drug resistance. As demonstrated in Fig. 3 and Fig. S2, our model allows simulation of how a population specialized to one environment could behave under another. Therefore, our framework may naturally extend to predict how a persister strategy optimal under one type of drug behaves under another. Our results already suggest new ways to design treatment strategies. Supplementary results (Fig. S6) suggest that, for all environments we model, persister production decreases monotonically with the growth rate. Temporarily exposing the cell colony to favorable conditions will decrease variable persister production, lowering the proportion of persisters, potentially making the population more susceptible to antibiotics. This is consistent with existing experimental observations where exposing a colony to a fresh growth medium before applying antibiotics decreases persistence (61), and is a simplistic example of drug sequencing.

We focus on a large, exponentially growing population in a homogeneous, temporally fluctuating, environment. Our framework is readily extensible to more complex growth models (such as logistic growth); heterogeneous populations containing more than two phenotypes; and the inclusion of demographic noise. We have not, however, considered spatial effects such as those corresponding to a spatially fluctuating environment or local crowding effects. A stochastic partial differential equation (SPDE) model would also allow for direct modelling of the chemical signals that regulate quorum sensing (62) allowing for population-level cooperation, however applying optimal control to these SPDE models is not straightforward. Further work is needed to elucidate the effect of demographic or sensor noise on a population’s ability to regulate the optimal persister proportion through subpopulation switching (23, 63). In large exponentially growing populations, we expect deterministic switching rates are an appropriate model of subpopulation switching. For small populations, our optimal control approach can be applied to study the effects of demographic noise by incorporating intrinsic noise into the stochastic growth model through the chemical Langevin equation.

Rapid evolution of cellular hedging strategies is experimentally reproducible (64): Van den Bergh *et al.* (14) found that exposing *E. coli* to daily antibiotic treatments increased survival by between three and 300-fold after just three treatments, and Rodriguez-Beltran *et al.* (65) study induced evolutionary changes in bacteria behaviour through repeated application of antibiotics. Evolutionary adaption of organisms such as bacteria within a *fitness landscape* is an active area of research (15), with an understanding that stochasticity plays a vital role (54). Financial mathematics techniques and cellular hedging can also be applied to study the evolutionary process as it occurs. For example, policy adjustment models quantify the cost of strategy change, and can be compared to mutation costs in a rapidly-evolving cell colony. Our framework can be directly applied in this context to model the adaption of a simple hedging strategy to an unfamiliar environment by modelling the time-derivative of the constant switching rates as a control.

Our analysis suggests important experimental avenues to further elucidate bacterial persistence. Alternating periods of growth and stress is still a common model of environmental volatility both (65) and in theoretical studies (9). We provide a new theoretical foundation for studying any Markovian model of environmental volatility. Supplementary results (Fig. S2) show a diversity in optimal responses when a strategy optimal under one type of environment reacts to another. An experimental study where bacteria are repeatedly exposed to a known volatile environment, potentially based upon the novel models of environmental volatility that we study, will provide insight into how bacteria adapt to a form of uncertainty that can be quantified. Our modelling framework can then predict both the emergent strategy, and how the population might behave when exposed to interventions such as antibiotics. Furthermore, sensitivity analysis on parameters in the environmental volatility model can aid experimental design by revealing what features of an environment have the largest effect on any optimal strategy. We expect, for instance, different responses to environments where changes happen continuously (for example, the Monod environment), compared to where changes are due to shocks (for example, the Poisson environment).

## Conclusion

We provide new insight into cellular hedging through an explicit connection between the study of biological population dynamics and the mathematics of financial risk management. We present a new stochastic model of bacteria growth in a volatile environment and apply optimal control theory to probe the persister strategy that maximizes the per-capita growth rate. A fundamental result of our study is to provide a solid theoretical understanding of why persistence is only advantageous in the presence of environmental volatility, demonstrating that the study of bacterial persistence must take a stochastic perspective.Our model of cellular hedging has clinical significance and can be applied to improve the efficacy of antimicrobial therapies. Furthermore, the framework we develop can be applied to more complex models of bacterial population dynamics, and offers an opportunity for future generalisations to explore cellular decision making in a broader context, including in cases where sensor or demographic noise are significant.Many seemingly complex cellular phenomena from bet-hedging in cancers (7, 67, 68), herpes viruses (69) and HIV (70) to decision making in the epithelial-mesenchymal transition (71) can be modelled using ideas from cellular hedging and by furthering the unification of mathematical finance and biology.

## Acknowledgements

This work is supported by the Australian Research Council (DP170100474) and the Air Force Office of Scientific Research (BAA-AFRL-AFOSR-2016-0007). We thank Jacob Scott and two anonymous referees for their helpful comments.

## Author Contributions

All authors designed the research; A.P.B. performed the research and wrote the manuscript; A.P.B., T.M., and J.A.S. implemented the numerical techniques. All authors provided direction, feedback and gave approval for final publication.

## Data availability

This study contains no experimental data. Code used to produce the numerical results is available on GitHub at github.com/ap-browning/persisters.

