## Supplementary Material for "Persistence as an optimal hedging strategy"

### Supplementary material for
### "Persistence as an optimal hedging strategy"

A.P. Browning*, J.A. Sharp, T. Mapder, C.M. Baker, K. Burrage and M.J. Simpson

**5 Optimal persister proportion** **16**

### 1 Additional environments

In the main document, we present results for three stochastic environments (Constant, Monod and Poisson). In this supporting material document, we present and produce results for two additional environments. These environments are shown in Fig. S1*a*–*e*. In Fig. S1 we reproduce results in Fig. 2 including these two additional environments.

4. *Ornstein-Uhlenbeck.* (Fig. S1*d*) By considering that the environment moves continuously around $\mu_t = \mu_G$ and always reverts to this value, we can model $\zeta_t$ as an Ornstein-Uhlenbeck process (1) with $m(\zeta) = \zeta$. In this case we have

$$\mathrm{d}\zeta_t = \gamma(\mu_G - \zeta_t)\,\mathrm{d}t + \kappa\,\mathrm{d}W_t^{(3)}\,, \tag{S1}$$

   where $\gamma$ and $\kappa$ represent the strength of the mean-reversion process and variability, respectively; and $\mathrm{d}W_t^{(3)}$ is a Wiener process independent to both $\mathrm{d}W_t^{(1)}$ and $\mathrm{d}W_t^{(2)}$ that satisfies $W_{t+h}^{(i)} - W_t^{(i)} \sim \mathcal{N}(0,h)$ where $\mathcal{N}(0,h)$ denotes a normal distribution with mean zero and variance $h$. For numerical results, we choose $\mu_G = 2\,\mathrm{h}^{-1}$, $\gamma = 1$ and $\kappa = 0.1$.

5. *Duffing.* (Fig. S1*e*) We can mimic the effects of the Poisson switching process using a Duffing oscillator. We consider that the environment, $\zeta_t$, switches randomly between two stable states at $\zeta_t = 0, 1$. These states are separated by an asymmetric instability, such that the state at $\zeta_t = 0$ is rarer than that at $\zeta_t = 1$. We describe this with the Duffing oscillator

$$\mathrm{d}\zeta_t = \gamma\zeta_t(1-\zeta_t)(\zeta_t - b)\,\mathrm{d}t + \kappa\,\mathrm{d}W_t^{(3)}\,, \tag{S2}$$

   where $\gamma$ is the strength of the oscillator; $b$ is the location of the instability, $\kappa$ represents the strength of the fluctuations of the oscillator; and $\mathrm{d}W_t^{(3)}$ is a Wiener process independent to both $\mathrm{d}W_t^{(1)}$ and $\mathrm{d}W_t^{(2)}$ that satisfies $W_{t+h}^{(i)} - W_t^{(i)} \sim \mathcal{N}(0,h)$ where $\mathcal{N}(0,h)$ denotes a normal distribution with mean zero and variance $h$.

   We couple the dependence of the growth rate to the environment by setting $m(\zeta_t) = \mu_G + (\mu_G - \mu_S)\zeta_t$ so that for $\zeta_t \approx 1$, $m(\zeta_t) \approx \mu_G$ and for $\zeta_t = 0$, $\mu \approx \mu_S$. We choose $\gamma = 20$, $b = 0.4$, $\kappa = 0.5$ and $\mu_S = -2\,\mathrm{h}^{-1}$.

In Fig. S2 we examine how a persister strategy that is optimal under one type of environment behaves under another, unfamiliar environment. We assume that cells monitor their growth rate, $\mu_t$, and couple the growth rate to the optimal control by considering $\phi_t^* = \phi^*(\theta_t, \zeta_t)$ where $\zeta_t = m^{-1}(\cdot)$ denotes the inverse of the growth rate coupling function $m(\zeta)$. In other words, the cells measure the current environment state using the growth rate. In Fig. S2*e*–*h* we show how a persister strategy that is optimal under the Monod environment behaves when exposed to a growth rate from: the Monod environment (Fig. S2*e*); the Poisson environment (Fig. S2*f*); the Ornstein-Uhlenbeck environment (Fig. S2*g*); and, the Duffing environment (Fig. S2*h*). In subsequent rows we repeat this for persister strategies optimal under: the Poisson environment (Fig. S2*i*–*l*); the Ornstein-Uhlenbeck environment (Fig. S2*m*–*p*); and, the Duffing environment (Fig. S2*q*–*t*).

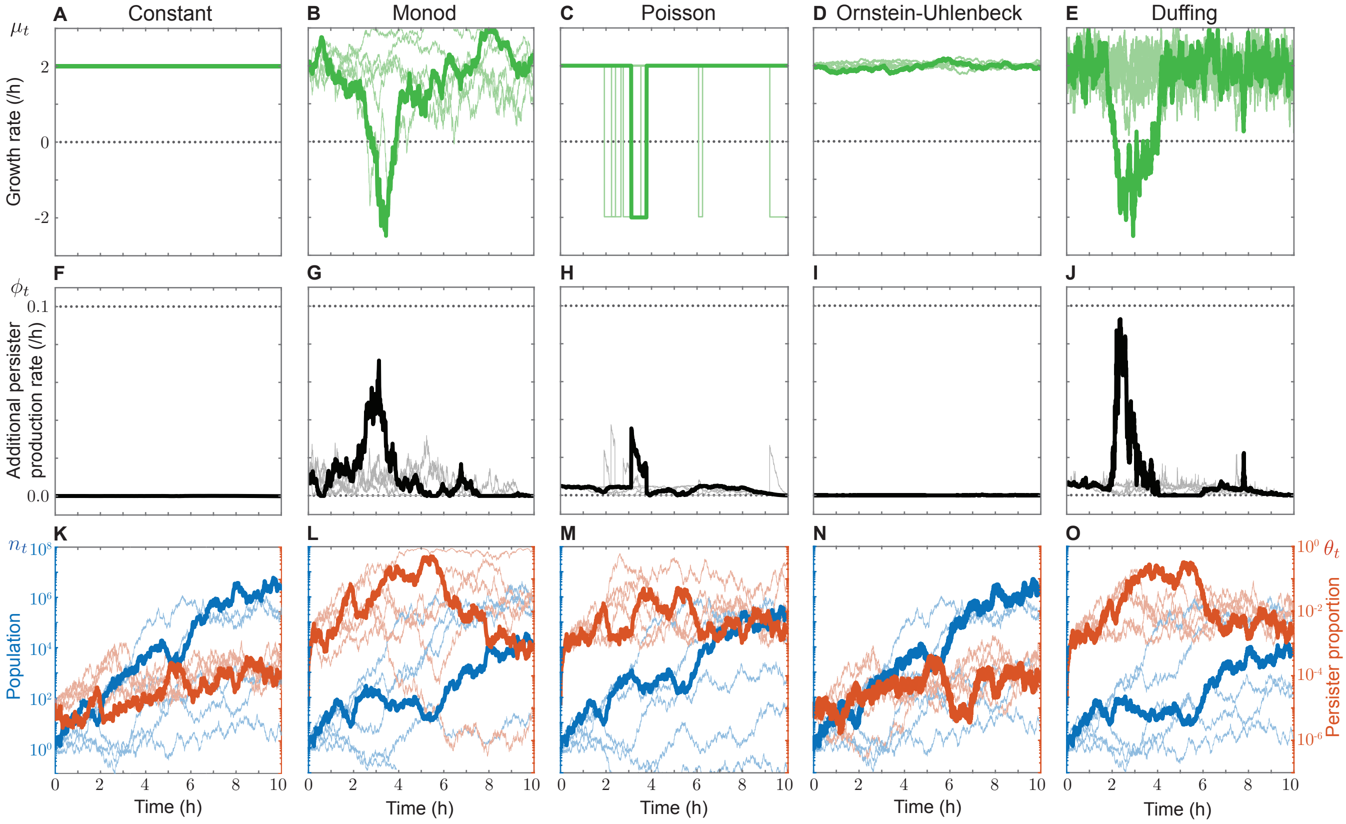

**Fig. S1.** Optimal persister production in cell populations under various types of environment showing: (a)-(e) the growth rates produced by each environment; (f)-(j) the persister production; and, (k)-(o) the population (blue, left scale) and persister proportion (red, right scale). The seeds used to generate the Wiener process were fixed for all environments. Also shown are five additional, independent, realisations (semi-transparent).

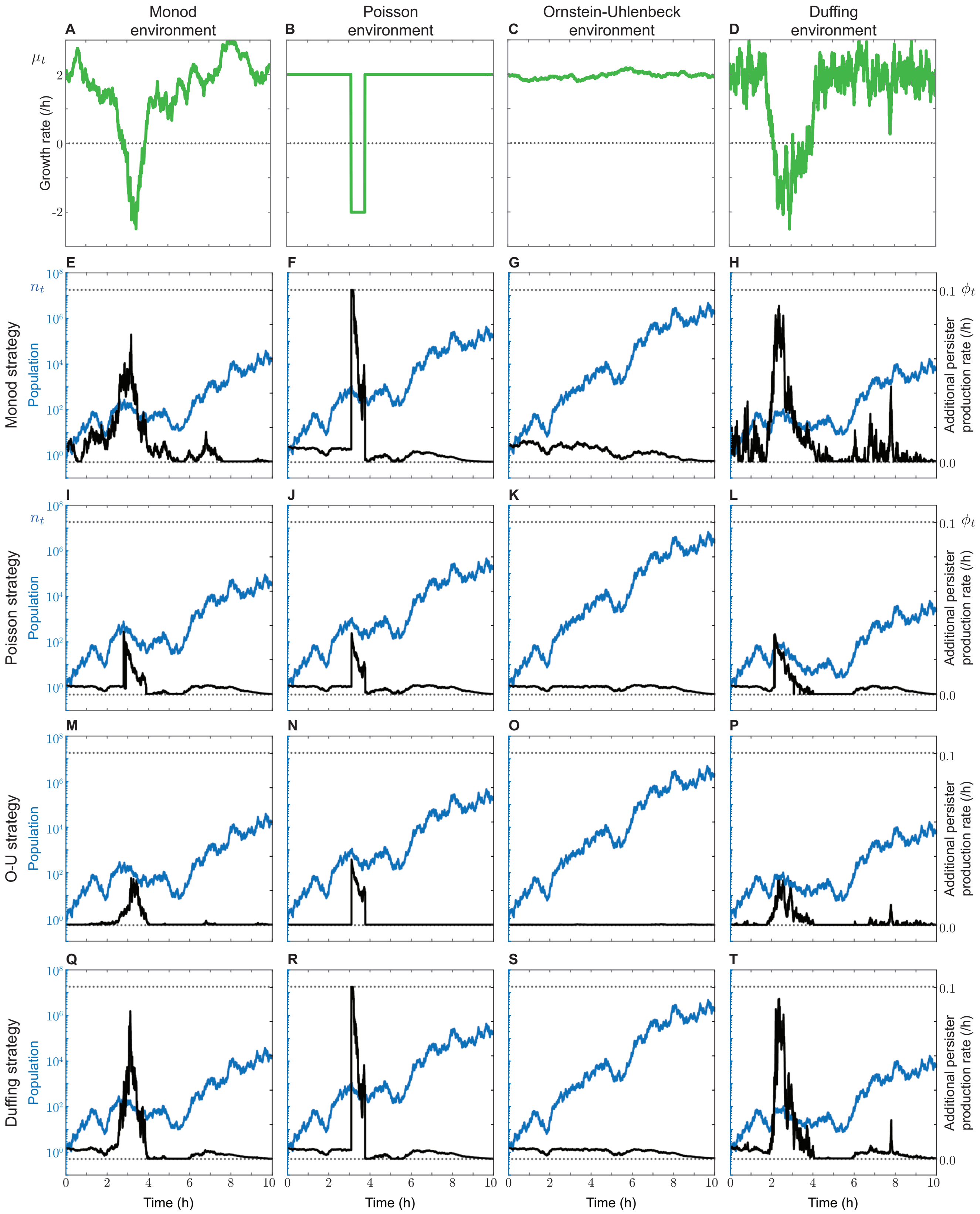

**Fig. S2.** Behaviour under each cell strategy in response to an unfamiliar environment. The rows correspond to each optimal cell strategy, the columns to each environment which the cells respond to. Note that (e), (j), (o) and (t) correspond to (g)–(j) of Fig. S1.

### 2 Constant persister production

Here, we assume that the cell population can use the switching rates, $u$ and $v$, to steer the steady state proportion, $\hat{\theta}_t$, based on an optimality assumption. We only consider this type of persistence in the presence of a constant environment in growth, so that $\mu_t = \mu_G$. Allowing $\hat{\theta}_t \in [0, 1]$ to be a control reduces the system to a one-dimensional control problem with state equation given by the Itô form

$$\frac{\mathrm{d}n_t}{n_t} = (1 - \hat{\theta}_t(1 - \varepsilon))\mu_G \,\mathrm{d}t + \sqrt{(1 - \hat{\theta}_t)^2\sigma^2 + \hat{\theta}_t^2\eta^2}\,\mathrm{d}W_t\,, \tag{S3}$$

where $\hat{\theta}_t$ is now a control. Setting $\mathbb{E}(\mathrm{d}\theta_t) = 0$ in Eq. 4b, reveals reveals the relationship between $u$, $v$ and $\hat{\theta}_t$:

$$\begin{aligned} 0 &= (1 - \theta_t)u - \theta_t v - (1 - \theta_t)\theta_t\big((1 - \varepsilon)\mu_t - \sigma^2(1 - \theta_t) + \eta^2\theta_t\big), \\ \Rightarrow u &= \hat{\theta}_t\left(\frac{v}{1 - \hat{\theta}_t} + \eta^2\hat{\theta}_t - (1 - \hat{\theta}_t)\sigma^2 + (1 - \varepsilon)\mu\right). \end{aligned}$$

As $\phi_t = 0$, the fitness measure (Eq. 7) reduces to

$$J_{\hat{\theta}} = \mathbb{E}\big[\log_{\mathrm{e}}(n_T)\big],$$

where $n_t$ is described by Eq. S3, and $n_T$ represents $n_t$ at terminal time, $t = T$.

We solve this optimal control problem using Hamilton-Jacobi-Bellman (HJB) optimal control theory (1, 2). We define the value function

$$V(x, s) = \max_{\hat{\theta}_t} \mathbb{E}\big[\log_{\mathrm{e}}(n_T)\big], \quad \text{where } n_s = x, \tag{S4}$$

and $n_t$ is described by Eq. S3. Therefore, $V$ describes the optimal payoff obtainable, starting at state $x$ at time $s$. HJB describes $V$ as the solution of a partial differential equation (PDE) (1, 2), given by

$$0 = \max_{\hat{\theta}_t}\left(\frac{\partial V}{\partial s} + (1 - \hat{\theta}_s(1 - \varepsilon))\mu_G x\frac{\partial V}{\partial x} + \frac{1}{2}x^2\left((1 - \hat{\theta}_t)^2\sigma^2 + \hat{\theta}_s^2\eta^2\right)\frac{\partial^2 V}{\partial x^2}\right), \quad V(x, T) = \log_{\mathrm{e}}(x). \tag{S5}$$

Here, the terminal condition of the PDE is obtained by substituting $s = T$ in Eq. S4.

As Eq. S5 is quadratic in $\hat{\theta}_t$, we can carry out the maximization by setting the derivative to zero to see that the optimal control, $\hat{\theta}_t^*$, is given by

$$\hat{\theta}_s^* = \frac{(1 - \varepsilon)\mu_G\dfrac{\partial V}{\partial x} + x\sigma^2\dfrac{\partial^2 V}{\partial x^2}}{x\left(\sigma^2 + \eta^2\right)\dfrac{\partial^2 V}{\partial x^2}}. \tag{S6}$$

Substituting Eq. S6 into Eq. S5 gives a nonlinear PDE that can be solved to give $V(x, s)$ and hence $\hat{\theta}_t^*$. Noting the form of the terminal condition, the ansatz

$$V(x, s) = h(s) + \log_{\mathrm{e}}(x), \quad h(T) = 0, \tag{S7}$$

is consistent with the PDE, with

$$h(s) = \exp\left(\frac{(T - s)\big\{\mu_G\big[(1 - \varepsilon)^2\mu_G + 2\varepsilon\sigma^2\big] + \eta^2(2\mu_G - \sigma^2)\big\}}{2(\sigma^2 + \eta^2)}\right).$$

To obtain an algebraic expression for the optimal control strategy, we can substitute the analytical solution for $V(x, s)$ (Eq. S7) into Eq. S6 to obtain an expression for $\hat{\theta}_t^*$. Noting that $\hat{\theta}_t^* \in [0, 1]$, we can write this as

$$\hat{\theta}_t^* = \max\left(0, \min\left(1, \frac{\sigma^2 - (1 - \varepsilon)\mu_G}{\eta^2 + \sigma^2}\right)\right). \tag{S8}$$

As Eq. S5 is quadratic in the control, we can verify the optimal solution is, indeed, a maximum by examining the sign of the coefficient of $\hat{\theta}_t^2$. We see that

$$\text{sign}\left(\frac{1}{2}x^2\eta^2\frac{\partial^2 V}{\partial x^2}\right) = -1,$$

so we always obtain a maximum.

### 3 Variable persister production

We first apply Itô's lemma* (1) to the change of variables,

$$\tilde{n}_t = \log(r_t + p_t), \quad \theta_t = \frac{p_t}{n_t},$$

giving

$$\mathrm{d}\begin{bmatrix}\tilde{n}_t\\ \theta_t\end{bmatrix} = \begin{bmatrix} f^{(\tilde{n})}(\theta_t, \zeta_t)\\ f^{(\theta)}(\theta_t, \zeta_t, \phi_t)\end{bmatrix}\mathrm{d}t + \boldsymbol{\Sigma}(\theta_t)\,\mathrm{d}\mathbf{W}_t\,, \tag{S9}$$

where $\mathrm{d}\mathbf{W}_t = [\mathrm{d}W_t^{(1)}\,, \mathrm{d}W_t^{(2)}]^{\mathrm{tr}}$ denotes a two-dimensional Wiener process with independent components, where superscript $^{\mathrm{tr}}$ the matrix transpose, and

$$\boldsymbol{\Sigma}(\theta_t)\boldsymbol{\Sigma}(\theta_t)^{\mathrm{tr}} = \begin{bmatrix}\Sigma^{(\tilde{n}\tilde{n})}(\theta_t) & \Sigma^{(\tilde{n}\theta)}(\theta_t)\\ \Sigma^{(\tilde{n}\theta)}(\theta_t) & \Sigma^{(\theta\theta)},(\theta_t)\end{bmatrix}, \tag{S10a}$$

$$f^{(\tilde{n})}(\theta_t, \zeta_t) = (1-(1-\varepsilon)\theta_t)m(\zeta_t) - \frac{\sigma^2(1-\theta_t)^2}{2} - \frac{\eta^2\theta_t^2}{2}, \tag{S10b}$$

$$f^{(\theta)}(\theta_t, \zeta_t, \phi_t) = (u+\phi_t)(1-\theta_t) - v\theta_t - (1-\theta_t)\theta_t(\eta^2\theta_t + m(\zeta_t)(1-\varepsilon) - (1-\theta_t)\sigma^2) \tag{S10c}$$

$$\Sigma^{(\tilde{n}\tilde{n})}(\theta_t) = \sigma^2(1-\theta_t)^2 + \eta^2\theta_t^2, \tag{S10d}$$

$$\Sigma^{(\theta\theta)}(\theta_t) = (\sigma^2+\eta^2)(1-\theta_t)^2\theta_t^2, \tag{S10e}$$

$$\Sigma^{(\tilde{n}\theta)}(\theta_t) = (1-\theta_t)\theta_t[\eta^2\theta_t - \sigma^2(1-\theta_t)]. \tag{S10f}$$

Here, $\zeta_t$ is described by a state equation that differs between environments and we express $\boldsymbol{\Sigma} = \boldsymbol{\Sigma}(\theta_t)$ to denote the dependence of, for example, $\boldsymbol{\Sigma}$ on only $\theta_t$.

**3.1. Continuous environments.** For each continuous environment, we can express the state equation for $\zeta_t$ as

$$\mathrm{d}\zeta_t = f^{(\zeta)}(\zeta_t)\,\mathrm{d}s + \sqrt{\Sigma^{(\zeta\zeta)}(\zeta_t)}\,\mathrm{d}W_t^{(3)}\,. \tag{S11}$$

Here, $\mathrm{d}W_t^{(3)}$ is a Wiener process scaled by an intensity $\sqrt{\Sigma^{(\zeta\zeta)}(\zeta_t)}$, and is independent of both $\mathrm{d}W_t^{(1)}$ and $\mathrm{d}W_t^{(2)}$

To find the optimal control, we define the value function, $V(x,y,z,s)$, such that

$$V(x,y,z,s) = \max_{\phi_t\in\mathcal{U}}\mathbb{E}\left[\int_s^T \alpha\phi_t^2\,\mathrm{d}s + x_T\right],\ (\tilde{n}_s, \theta_s, \zeta_s) = (x,y,z), \tag{S12}$$

where $\phi_t$ is the control.

HJB describes $V$ as the solution of PDE (1, 3), given by

$$0 = \max_{\phi_s\in\mathcal{U}}\left(\mathcal{A}V + \alpha\phi_t^2\right) = \max_{\phi_t\in\mathcal{U}}\mathcal{H}$$

where $\mathcal{A}V$ is the infinitesimal generator of $V$ with respect to the stochastic process given by corresponding state equations, and we have defined $\mathcal{H} = \mathcal{A}V + \alpha\phi_t^2$ for notational clarity. For the the continuous environment, the

*__Itô's lemma.__ For the stochastic process given by $\mathrm{d}\mathbf{X}_t = \mathbf{M}(\mathbf{X}_t)\,\mathrm{d}t + \mathbf{S}(\mathbf{X}_t)\,\mathrm{d}\mathbf{W}_t$ — where $\mathbf{X}_t \in \mathbb{R}^2$, $\mathbf{M} : \mathbb{R}^2 \to \mathbb{R}^2$, $\mathbf{S} : \mathbb{R}^2 \to \mathbb{R}^{2\times 2}$ and $\mathbf{W}_t$ is a two-dimensional Wiener process — then Itô's lemma states that

$$\mathrm{d}f(\mathbf{X}_t) = \left[(\nabla_{\mathbf{X}} f)^{\mathrm{tr}}\,\mathbf{M}(\mathbf{X}_t) + \frac{1}{2}\mathrm{Trace}(\mathbf{S}(\mathbf{X}_t)^{\mathrm{tr}}\,(H_{\mathbf{X}}f)\mathbf{S}(\mathbf{X}_t))\right]\mathrm{d}t + (\nabla_{\mathbf{X}} f)^{\mathrm{tr}}\,\mathbf{S}(\mathbf{X}_t)\,\mathrm{d}\mathbf{W}_t\,.$$

Here, $H_{\mathbf{X}}f$ denotes the Hessian matrix for the function $f$; superscript $^{\mathrm{tr}}$ the matrix transpose; and Trace$(\cdot)$ the matrix trace. In our study, the untransformed variables are denoted $\mathbf{X} = [r_t, p_t]^{\mathrm{tr}}$ and the transformed system denoted $\mathbf{Y} = [f(r_t,p_t), g(r_t,p_t)]^{\mathrm{tr}} = [\tilde{n}_t, \theta_t]^{\mathrm{tr}}$.

state equations are given by Eqs. S9 and S11 so $V$ is described by the PDE

$$0 = \max_{\phi_t \in \mathcal{U}} \left\{ \frac{\partial V}{\partial s} + f^{(\tilde{n})}\frac{\partial V}{\partial x} + f^{(\theta)}\frac{\partial V}{\partial y} + f^{(\zeta)}\frac{\partial V}{\partial z} + \frac{\Sigma^{(\tilde{n}\tilde{n})}}{2}\frac{\partial^2 V}{\partial x^2} + \frac{\Sigma^{(\theta\theta)}}{2}\frac{\partial^2 V}{\partial y^2} + \frac{\Sigma^{(\zeta\zeta)}}{2}\frac{\partial^2 V}{\partial z^2} + \Sigma^{(\tilde{n}\theta)}\frac{\partial^2 V}{\partial x \partial y} + \alpha\phi_t^2 \right\},$$
$$V(x,y,z,T) = x.$$

As all of the coefficients, $f^{(\cdot)}$ and $\Sigma^{(\cdot\cdot)}$, do not depend on $x$, we introduce the ansatz $V(x,y,z,s) = \Psi(y,z,s)+x$, leading to a two-dimensional PDE

$$0 = \max_{\phi_t \in \mathcal{U}} \left\{ \frac{\partial \Psi}{\partial s} + f^{(\tilde{n})} + f^{(\theta)}\frac{\partial \Psi}{\partial y} + f^{(\zeta)}\frac{\partial \Psi}{\partial z} + \frac{\Sigma^{(\theta\theta)}}{2}\frac{\partial^2 \Psi}{\partial y^2} + \frac{\Sigma^{(\zeta\zeta)}}{2}\frac{\partial^2 \Psi}{\partial z^2} + \alpha\phi_t^2 \right\}, \quad \Psi(y,z,T) = 0. \tag{S13}$$

Note that in the main document, we do not apply the transformation $x \to \log_e(x)$, so the ansatz is equivalent to $V(x,y,z,s) = \Psi(y,z,s) + \log_e(x)$.

For brevity, we express $\mathcal{H}$ as

$$\mathcal{H} = \mathcal{H}_\phi(y,\phi_t,s) + \mathcal{H}_\star(y,z,s),$$

so that

$$\operatorname*{argmax}_{\phi_t \in \mathcal{U}} \mathcal{H} = \operatorname*{argmax}_{\phi_t \in \mathcal{U}} \mathcal{H}_\phi = \phi_t^*,$$

and

$$\mathcal{H}_\phi = \alpha\phi_t^2 + \frac{\partial V}{\partial y}(1-y)\phi_t,$$

for all continuous environments. This gives the optimal control $\phi_t^* \in [0, 0.1]$,

$$\phi_t^* = \max\left[0, \min\left[0.1, -\frac{1}{2\alpha}\frac{\partial V}{\partial y}(1-y)\right]\right]. \tag{S14}$$

It is evident that $\partial^2\mathcal{H}_\phi/\partial\phi^2 = 2\alpha < 0$ (since $\alpha < 0$), so we always obtain a maximum.

**3.2. Poisson environment.** We can replace the variable rate Poisson process, $\mathrm{d}P_t$ (described by Eq. 6a) with

$$\mathrm{d}\zeta_t = \rho(\zeta_t)\,\mathrm{d}P_t = \rho_1(\zeta_t)\,\mathrm{d}P_t^{(1)} + \rho_2(\zeta_t)\,\mathrm{d}P_t^{(2)}, \tag{S15}$$

where $\mathrm{d}P_t^{(1)}$ and $\mathrm{d}P_t^{(2)}$ have constant rates $\lambda_1 = 1/\tau_G$ and $\lambda_2 = 1/\tau_S$, respectively, and jump distances

$$\rho_1(\zeta_t) = -(\zeta_t+1) = \begin{cases} -2, & \zeta_t = +1, \\ 0, & \zeta_t = -1, \end{cases} \quad \rho_2(\zeta_t) = -(\zeta_t-1) = \begin{cases} 0, & \zeta_t = +1, \\ +2, & \zeta_t = -1. \end{cases}$$

For the the Poisson environment, the state equations are given by Eqs. S9 and S15 so $V$ is described by the PDE

$$0 = \max_{\phi_t \in \mathcal{U}} \left\{ \frac{\partial V}{\partial s} + f^{(\tilde{n})}\frac{\partial V}{\partial x} + f^{(\theta)}\frac{\partial V}{\partial y} + \frac{\Sigma^{(\tilde{n}\tilde{n})}}{2}\frac{\partial^2 V}{\partial x^2} + \frac{\Sigma^{(\theta\theta)}}{2}\frac{\partial^2 V}{\partial y^2} + \lambda_1\left[V\big|_{z=-1} - V + (z+1)\frac{\partial V}{\partial z}\right] \right.$$
$$\left. + \lambda_2\left[V\big|_{z=+1} - V + (z-1)\frac{\partial V}{\partial z}\right] + \alpha\phi_t^2 \right\}, \quad V(x,y,z,T) = x.$$

Introducing the ansatz $V(x,y,z,t) = \Psi(y,z,t) + x$; and, since $m(\zeta_t) = \mathrm{sign}(\zeta_t)$ so that $\partial V/\partial z = 0$, we have that

$$0 = \max_{\phi_t \in \mathcal{U}} \left\{ \frac{\partial \Psi}{\partial s} + f^{(\tilde{n})} + f^{(\theta)}\frac{\partial \Psi}{\partial y} + \frac{\Sigma^{(\theta\theta)}}{2}\frac{\partial^2 \Psi}{\partial y^2} + \lambda_1\left[\Psi\big|_{z=-1} - \Psi\right] + \lambda_2\left[\Psi\big|_{z=+1} - \Psi\right] + \alpha\phi_t^2 \right\}, \quad \Psi(y,z,T) = 0. \tag{S16}$$

As Eq. S16 and the state equations only depend on $z$ spatially at $z = \pm 1$, we write Eq. S16 as a coupled system of PDEs where $\Psi^+$ denotes $\Psi$ at $z = 1$; and $\Psi^-$ denotes $\Psi$ at $z = -1$,

$$0 = \frac{\partial \Psi^+}{\partial s} + f^{(\tilde{n})}(y, +1) + f^{(\theta)}(y, +1, \phi_t^*) \frac{\partial \Psi^+}{\partial y} + \frac{\Sigma^{(\theta\theta)}(y)}{2} \frac{\partial^2 \Psi^+}{\partial y^2} + \lambda_1 \left[ \Psi^- - \Psi^+ \right] + (\phi_t^*)^2 \alpha, \tag{S17a}$$

$$0 = \frac{\partial \Psi^-}{\partial s} + f^{(\tilde{n})}(y, -1) + f^{(\theta)}(y, -1, \phi_t^*) \frac{\partial \Psi^-}{\partial y} + \frac{\Sigma^{(\theta\theta)}(y)}{2} \frac{\partial^2 \Psi^-}{\partial y^2} + \lambda_2 \left[ \Psi^+ - \Psi^- \right] + (\phi_t^*)^2 \alpha. \tag{S17b}$$

Here, the optimal control, $\phi_t^*$ is given by Eq. S14.

### 4 Numerical techniques

To solve the HJB equation (Eq. S13) we employ a numerical scheme with a logarithmically spaced grid in $y$ near $y = 0$ and $y = 1$ (Fig. S3$a$), and constant spacing in $z$. We use up-winding to approximate derivatives in the $y$ direction (Fig. S3$b$). At the boundaries, we note that $\Sigma^{(\theta\theta)}(0, z) = \Sigma^{(\theta\theta)}(1, z) = 0$, so we only need to approximate the first derivative on the boundaries in $y$. For all other boundaries, we linearly extrapolate the two nearest nodes to estimate the derivatives on the boundary (Fig. S3$c$). We use the explicit Euler method with $\Delta t = 1/30000$ to integrate backwards time from the terminal condition at $t = T$ to $t = 0$. To verify the accuracy of our solution, we compare the integral of the solution with both $\Delta t = 1/30000$ and $\Delta t = 1/60000$, and find a negligible difference. We show the numerical solutions at $t = 0$ (the PDE is solved backwards in time from $t = T$), for each environment, in Fig. S4.

We use the Euler-Maruyama algorithm to simulate the SDEs forward. In solving the HJB equation, we store the discretized solution for the control, and interpolate using `interpn` in MATLAB (4) to get the control. In Eq. S5 we verify the accuracy of our numerical methods. The numerical solution to the HJB equation at $t = 0$ gives, by definition (Eq. S12),

$$V(x, y, z, 0) = \mathbb{E}\left[\int_0^T \alpha\phi_t^2 \, \mathrm{d}t + \log_{\mathrm{e}}(n_T)\right], \tag{S18}$$

which we can compare to the ensemble average of 5000 realizations of the SDE. In Eq. S5 we show both our numerical solution and the ensemble average at $\mu_0 = 2$ and $n_0 = 1$.

In the following subsections we provide full details of our numerical solutions. Code used to produce the numerical results is available on GitHub at `github.com/ap-browning/persisters`.

**4.1. Spatial discretization.** We note that $y \in [0, 1]$ and $z \in \mathbb{R}$, however typically we find that $z \sim \mathcal{O}(1)$ in realizations of the SDE for all the environments considered. We denote the discretized domain as $y_i$ and $z_j$, so that our finite difference approximation of the solution at each grid point, $\Psi(y_i, z_j, s)$, is denoted by

$$\Psi(y_i, z_j, s) \approx \overline{\Psi}_{i,j}(s),\ i \in \mathcal{I} = \{1, 2, ..., I\},\ j \in \mathcal{J} = \{1, 2, ..., J\}.$$

*Discretization in y.* We apply apply variable grid spacing in $y$, demonstrated in Fig. S3$a$,$b$. We set $y_1 = 0$ and $y_I = 1$. For $10^{-5} \leq y \leq 0.1$ and $0.9 \leq y \leq 1 - 10^{-5}$ we apply logarithmic grid spacing, such that $y_{50} = 0.1$ and $y_{81} = 0.9$. For $0.1 < y < 0.9$, we apply constant grid spacing. In this work, we find $I = 130$ is sufficient under visual inspection.

*Discretization in z.* We apply constant grid spacing in $z$ such that $z_j - z_{j-1} = \Delta z \;\forall\, j$. We solve over a sufficiently large domain in the $z$ direction to contain all realizations of corresponding SDE. In this work, we find that $J = 100$ is sufficient under visual inspection, except the Ornstein-Uhlenbeck environment where we set $J = 200$. The discretization in $z$ is shown in Fig. S3$b$.

**4.2. Derivatives.** We approximate derivatives in for central nodes using standard techniques, that we detail here. For the boundaries, we linearly extrapolate using the two nearest nodes, as shown in Fig. S3$c$.

*Derivatives in y.* We apply an up-winding scheme for the first derivative in $y$. Accounting for the variable mesh in the $y$ direction, on the interior nodes we define

$$\frac{\partial\overline{\Psi}_{i,j}}{\partial y}^{\text{forw}} = \frac{\overline{\Psi}_{i+1,j}(s) - \overline{\Psi}_{i,j}(s)}{y_{i+1} - y_i},$$
$$\frac{\partial\overline{\Psi}_{i,j}}{\partial y}^{\text{back}} = \frac{\overline{\Psi}_{i,j}(s) - \overline{\Psi}_{i-1,j}(s)}{y_i - y_{i-1}},$$

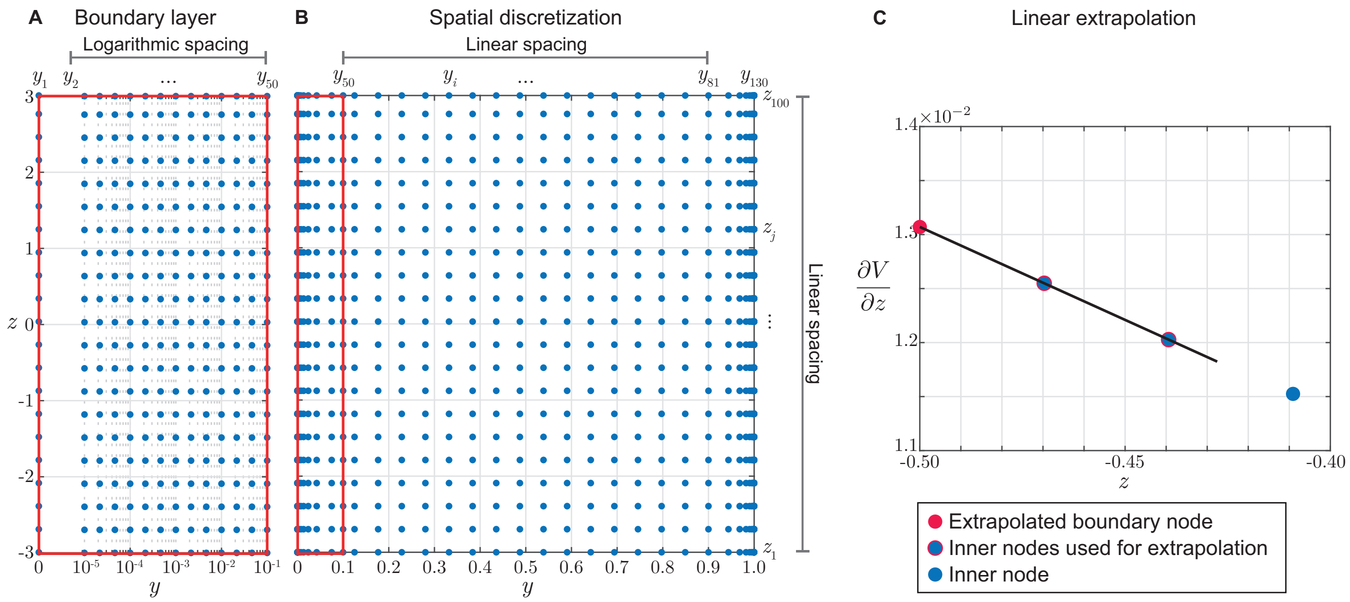

**Fig. S3.** Numerical scheme details. (a) and (b) demonstrate the spatial discretization. (a) is an inset of (b) (indicated in red) showing the logarithmic spacing for $y \leq 0.1$. (c) Linear extrapolation used to approximate the derivatives on the boundaries. A linear function is fit to the nearest two inner nodes (blue circles outlined in red) to extrapolate the value of the derivative on the boundary (red circle).

so that our finite difference approximation for the first derivative in $y$ is given by

$$\frac{\partial \Psi(y_i, z_i, s)}{\partial y} \approx \frac{\partial \overline{\Psi}_{i,j}}{\partial y} = \mathbb{1}_{\mathbb{R}^+}\left(f^{(\theta)}(y_i, z_j, (\overline{\phi_t^*})_{i,j})\right) \frac{\partial \overline{\Psi}_{i,j}}{\partial y}^{\text{forw}} + \mathbb{1}_{\mathbb{R}^-}\left(f^{(\theta)}(y_i, z_j, (\overline{\phi_t^*})_{i,j})\right) \frac{\partial \overline{\Psi}_{i,j}}{\partial y}^{\text{back}}. \tag{S19}$$

Here, $\mathbb{1}_{\mathbb{R}^+}(x)$ is an indicator function that takes the value 1 if $x \in \mathbb{R}^+$, and 0 otherwise. Eq. S19 applies for $i \in \mathcal{I} \backslash \{1, I\}$ and $j \in \mathcal{J}$. On the boundaries, where $i = 1$ and $i = I$, we linearly extrapolate using the nearest two nodes (Fig. S3$c$).

We apply a central difference for the second derivative in $y$, such that

$$\frac{\partial^2 \Psi(y_i, z_i, s)}{\partial y^2} \approx \frac{\partial^2 \overline{\Psi}_{i,j}}{\partial y^2} = \frac{2(y_i - y_{i-1})(\overline{\Psi}_{i+1,j}(s) - \overline{\Psi}_{i,j}(s)) - (y_{i+1} - y_i)(\overline{\Psi}_{i,j}(s) - \overline{\Psi}_{i-1,j}(s))}{(y_{i+1} - y_{i-1})(y_i - y_{i-1})(y_{i+1} - y_i)}, \tag{S20}$$

Eq. S20 applies for $i \in \mathcal{I} \backslash \{1, I\}$ and $j \in \mathcal{J}$. For $i = 1$ and $i = I$, the the coefficient of $\partial^2 \Psi / \partial y^2$ in the PDE, $\Sigma^{(\theta\theta)}(y_0) = \Sigma^{(\theta\theta)}(y_I) = 0$, so we do not need to extrapolate the second derivative in $y$.

*Derivatives in $z$.* On the interior nodes, we use central differences to approximate the first and second derivative in $z$, such that

$$\frac{\partial \Psi(y_i, z_i, s)}{\partial z} \approx \frac{\partial \overline{\Psi}_{i,j}}{\partial z} = \frac{\overline{\Psi}_{i,j+1}(s) - \overline{\Psi}_{i,j-1}(s)}{2\Delta z}, \tag{S21}$$

$$\frac{\partial^2 \Psi(y_i, z_i, s)}{\partial z^2} \approx \frac{\partial^2 \overline{\Psi}_{i,j}}{\partial z^2} = \frac{\overline{\Psi}_{i,j+1}(s) - 2\overline{\Psi}_{i,j}(s) + \Psi_{i,j-1}}{\Delta z^2}. \tag{S22}$$

Eqs. S21 and S22 apply for $i \in \mathcal{I}$ and $j \in \mathcal{J} \backslash \{1, J\}$. For $j = 1$ and $j = J$, we linearly extrapolate using the nearest two nodes (Fig. S3$c$).

**4.3. Time integration.** We apply the first-order explicit Euler method to integrate in time. Whereas the continuous environments involve solving a PDE for $\Psi(y, z, s)$, the Poisson environment involves solving the system of PDEs given by $\{\Psi^+(y, s), \Psi^-(y, s)\}$. Here, we describe our time-integration implementation separately for each type of PDE.

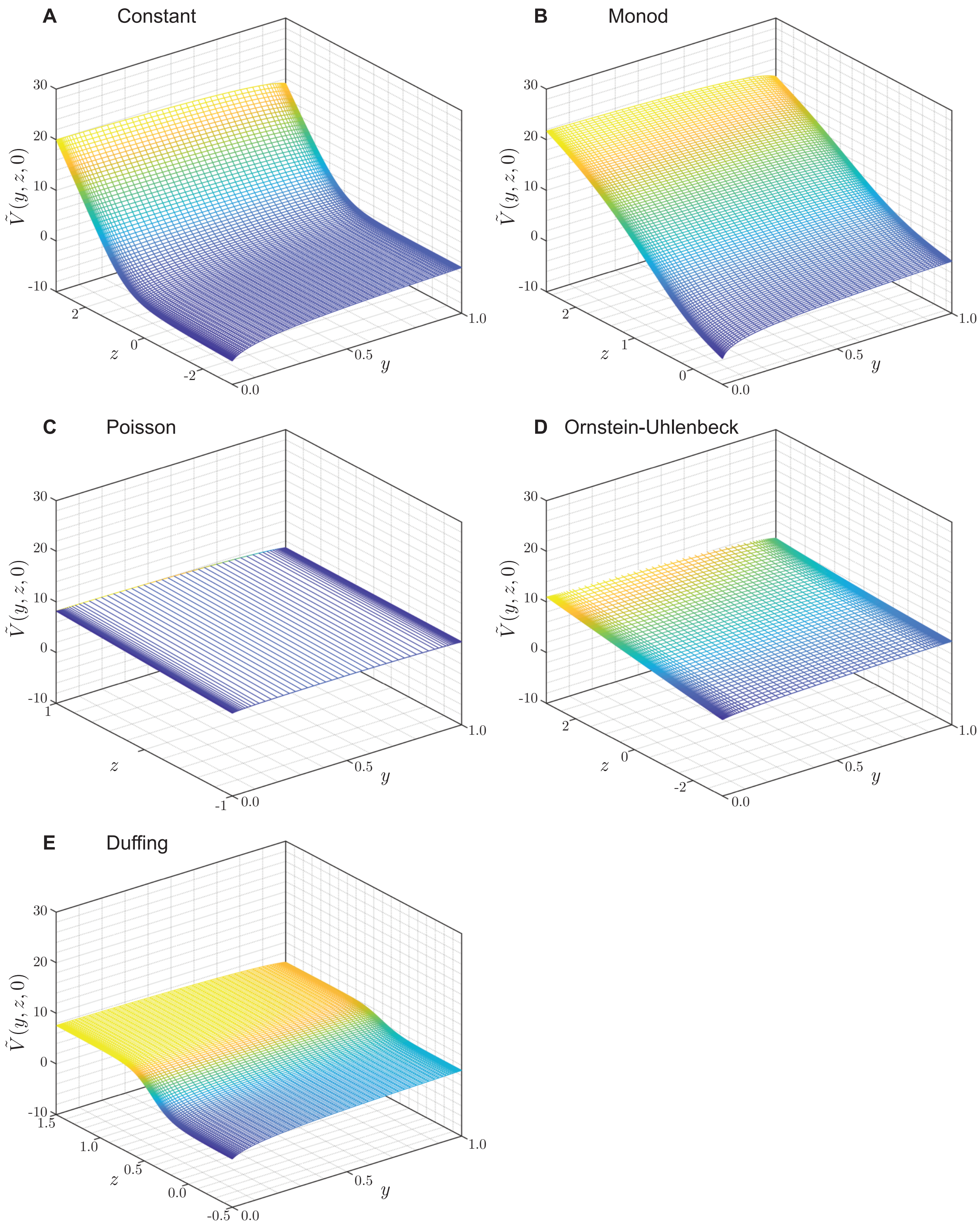

**Fig. S4.** Numerical solution to the HJB equation. Solution to the HJB equations, for each environment, at $t = 0\,\mathrm{h}$ (the HJB equations are solved backwards in time from $T = 10\,\mathrm{h}$), given by $\Psi(y, z, 0)$. In each case, $y = \theta_t$ and $z = \zeta_t$. The mesh for the Poisson environment only has two grid points in the $z$ direction, as the system corresponds to a system of two PDEs for $z \geq 0$ and $z < 0$.

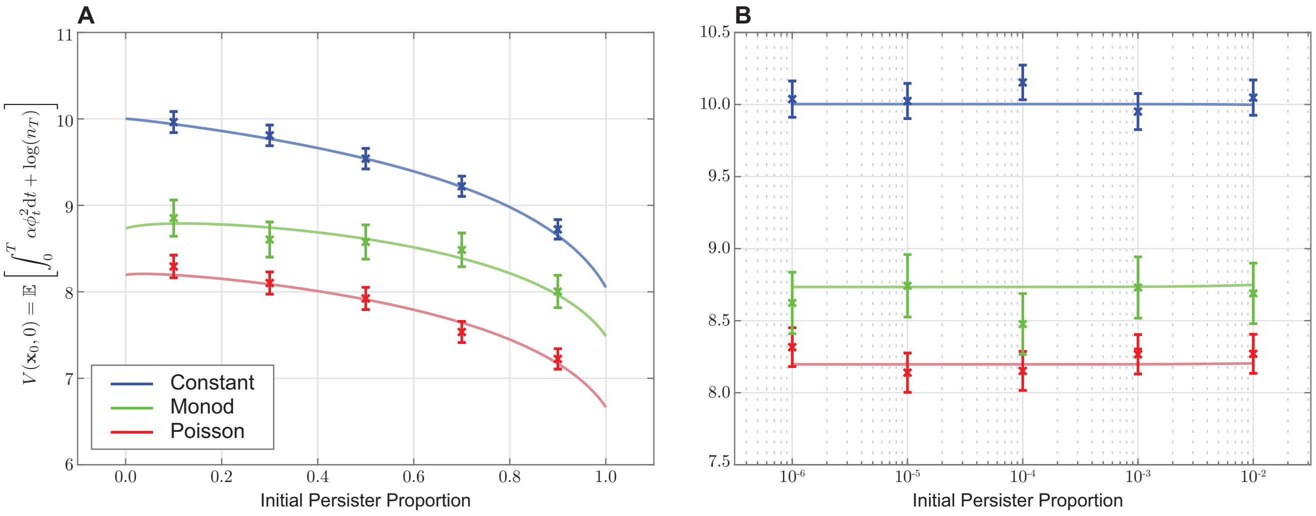

**Fig. S5.** Verification of numerical solution to the HJB equations. For each environment in the main document, we show the expected payoff, $V(\mathbf{x}_0, 0)$ (solid curve) as a function of the initial persister proportion, for $\mu_0 = 2\,\mathrm{h}^{-1}$ and $n_0 = 1$. In (a), we show the initial persister proportion in the linear scale; and in (b), we show the initial persister proportion in the log scale. We verify this numerical solution by showing the mean payoff from 5000 realizations of the SDE (crosses) along with a 95% confidence interval, assuming the payoff is normally distributed.

*Continuous environments.* Following the substitution of the optimal control (Eq. S14), we may express the PDE (Eq. S13) as

$$\frac{\partial \Psi}{\partial s} = F\left(s, y, z, \phi_t^*, \frac{\partial \Psi}{\partial y}, \frac{\partial \Psi}{\partial z}, \frac{\partial^2 \Psi}{\partial y^2}, \frac{\partial^2 \Psi}{\partial z^2}\right), \tag{S23}$$

where we define

$$F := -f^{(\tilde{n})}(y,z) - f^{(\theta)}(y,z,\phi_t^*)\frac{\partial \Psi}{\partial y} - f^{(\zeta)}(z)\frac{\partial \Psi}{\partial z} - \frac{\Sigma^{(\theta\theta)}(y)}{2}\frac{\partial^2 \Psi}{\partial y^2} - \frac{\Sigma^{(\zeta\zeta)}(z)}{2}\frac{\partial^2 \Psi}{\partial z^2} - \alpha \phi_t^2.$$

Applying the spatial discretization, we may write Eq. S23 as the system of ordinary differential equations (ODEs),

$$\frac{\mathrm{d}}{\mathrm{d}s}\overline{\Psi}_{i,j}(s) = \overline{F}_{i,j}(s) := F\left(s, y_i, z_i, (\overline{\phi_t^*})_{i,j}, \frac{\partial \overline{\Psi}_{i,j}}{\partial y}, \frac{\partial \overline{\Psi}_{i,j}}{\partial z}, \frac{\partial^2 \overline{\Psi}_{i,j}}{\partial y^2}, \frac{\partial^2 \overline{\Psi}_{i,j}}{\partial z^2}\right) \tag{S24}$$

where

$$(\overline{\phi_t^*})_{i,j} = \max\left(0, -\frac{1}{2\alpha}\frac{\partial \overline{\Psi}_{i,j}}{\partial y}\right),$$

which corresponds to the discretization of Eq. S14.

The ODE problem (Eq. S24) is coupled to a terminal condition at $s = T$. Therefore, we must integrate backwards in time to $s = 0$. To integrate from $s$ to $s - \Delta s$, we apply the explicit Euler scheme, so that

$$\begin{aligned}\overline{\Psi}_{i,j}(s - \delta s) &= \overline{\Psi}_{i,j}(s) + \int_s^{s-\Delta s} \overline{F}_{i,j}(\tau)\,\mathrm{d}\tau\,,\\ &\approx \overline{\Psi}_{i,j}(s) - \Delta s\, \overline{F}_{i,j}(s).\end{aligned}$$

*Poisson environment.* The system of PDEs for the Poisson environment (Eqs. S17) do not depend on $z$, and so we only discretize the domain in the $y$ direction. We define $\mathbf{\Psi} = [\Psi^+, \Psi^-]^{\mathrm{tr}}$ for brevity. Following the substitution of the optimal control (Eq. S14), we may express the system (Eqs. S17) as

$$\frac{\partial \mathbf{\Psi}}{\partial s} = \mathbf{F}\left(s, y, \phi_t^*, \Psi^+, \Psi^-, \frac{\partial \Psi^+}{\partial y}, \frac{\partial \Psi^-}{\partial y}, \frac{\partial^2 \Psi^+}{\partial y^2}, \frac{\partial^2 \Psi^-}{\partial y^2}\right), \tag{S25}$$

where we define

$$\mathbf{F} := \begin{bmatrix} -f^{(\tilde{n})}(y,+1) - f^{(\theta)}(y,+1,\phi_t^*)\dfrac{\partial \Psi^+}{\partial y} - \dfrac{\Sigma^{(\theta\theta)}(y)}{2}\dfrac{\partial^2 \Psi^+}{\partial y^2} - \lambda_1\left[\Psi^- - \Psi^+\right] - (\phi_t^*)^2\alpha, \\ -f^{(\tilde{n})}(y,-1) - f^{(\theta)}(y,-1,\phi_t^*)\dfrac{\partial \Psi^-}{\partial y} - \dfrac{\Sigma^{(\theta\theta)}(y)}{2}\dfrac{\partial^2 \Psi^-}{\partial y^2} - \lambda_2\left[\Psi^+ - \Psi^-\right] - (\phi_t^*)^2\alpha. \end{bmatrix} \tag{S26}$$

Applying the spatial discretization, we may write Eq. 4.3 as the system of ordinary differential equations (ODEs),

$$\frac{\mathrm{d}}{\mathrm{d}s}\overline{\mathbf{\Psi}}_i(s) = \overline{\mathbf{F}}_i(s) := \mathbf{F}\left(s, y_i, (\overline{\phi_t^*})_i, \overline{\Psi^+}_i, \overline{\Psi^-}_i, \frac{\partial \overline{\Psi^+}_i}{\partial y}, \frac{\partial \overline{\Psi^-}_i}{\partial y}, \frac{\partial^2 \overline{\Psi^+}_i}{\partial y^2}, \frac{\partial^2 \overline{\Psi^-}_i}{\partial y^2}\right) \tag{S27}$$

where

$$(\overline{\phi_t^*})_i = \max\left(0, -\frac{1}{2\alpha}\frac{\partial \overline{\Psi}_i}{\partial y}\right),$$

which corresponds to the discretization of Eq. S14.

The ODE problem (Eq. S24) is coupled to a terminal condition at $s = T$. Therefore, we must integrate backwards in time to $s = 0$. To integrate from $s$ to $s - \Delta s$, we apply the explicit Euler scheme, so that

$$\begin{aligned}\overline{\mathbf{\Psi}}_i(s - \delta s) &= \overline{\mathbf{\Psi}}_i(s) + \int_s^{s-\Delta s} \overline{\mathbf{F}}_i(\tau)\, \mathrm{d}\tau\,, \\ &\approx \overline{\mathbf{\Psi}}_i(s) - \Delta s\, \overline{\mathbf{F}}_i(s).\end{aligned}$$

**4.4. Solving the controlled SDE.** We apply the Euler-Maruyama algorithm (5) to solve the SDE with the optimal control. The numerical algorithm described above provides a discretized optimal control,

$$(\overline{\phi_t^*})_{i,j} = \overline{\phi_t^*}(y_i, z_i) \approx \phi_t^*(y_i, z_i).$$

To approximate $\phi_t^*(y, z)$ we interpolate the discretized function using `interpn` in MATLAB (4).

For all environments, the state equations for $\tilde{n}_t$ and $\theta_t$ are given by

$$\mathrm{d}\begin{bmatrix}\tilde{n}_t \\ \theta_t\end{bmatrix} = \mathbf{f}\big(\theta_t, \zeta_t, \phi_t^*(\theta_t, \zeta_t)\big)\, \mathrm{d}t + \mathbf{\Sigma}(\theta_t)\, \mathrm{d}\mathbf{W}_t\,,$$

where

$$\mathbf{f} := \begin{bmatrix} f^{(\tilde{n})}(\theta_t, \zeta_t) \\ f^{(\theta)}\big(\theta_t, \zeta_t, \phi_t^*(\theta_t, \zeta_t)\big)\end{bmatrix}$$

and $f^{(\tilde{n})}$, $f^{(\tilde{\theta})}$ and $\mathbf{\Sigma}(\theta_t)$ are given in Eq. S10.

For all environments, we initialize the numerical results by setting $\tilde{n}_0 = 0$, $\theta_0 = 1.19 \times 10^{-5}$ and choose $\zeta_0$ such that $\mu_t = m(\zeta_0) = 2$. To integrate $\tilde{n}_t$ and $\theta_t$ from $t$ to $t + \Delta t$, we use the Euler-Maruyama algorithm, such that

$$\begin{bmatrix}\tilde{n}_{t+\Delta t} \\ \theta_{t+\Delta t}\end{bmatrix} = \mathbf{f}\big(\theta_t, \zeta_t, \phi_t^*(\theta_t, \zeta_t)\big)\Delta t + \mathbf{\Sigma}(\theta_t)\sqrt{\Delta t}\begin{bmatrix}\Delta W_t^{(1)} \\ \Delta W_t^{(2)}\end{bmatrix}.$$

Here, $\Delta W_t^{(i)} \sim \mathcal{N}(0, 1)$ for $i = 1, 2$. In our study, we fix $\Delta t = 0.002$.

*Continuous environments.* To integrate $\zeta_t$ from $t$ to $t + \Delta t$, we use the Euler-Maruyama algorithm, such that

$$\zeta_{t+\Delta t} = f^{(\zeta)}(\zeta_t)\Delta t + \sqrt{\Sigma^{(\zeta\zeta)}(\zeta_t)\Delta t}\Delta W_t^{(3)}.$$

Here, $\Delta W_t^{(3)} \sim \mathcal{N}(0, 1)$.

*Poisson environment.* Here, we integrate $\zeta_t$ from $t$ to $t + \Delta t$ by considering

$$\zeta_{t+\Delta t} = \rho(\zeta_t)\Delta P_t,$$

where $\Delta P_t \sim \mathrm{Po}(\Delta t \lambda(\zeta_t))$. Here, $\rho(\zeta_t)$ and $\lambda(\zeta_t)$ are given in Eq. 6a.

### 5 Optimal persister proportion

The analytic expression for the optimal variable persister production (Eq. S14) suggests that the population uses $\phi_t^*$ to steer the population toward the optimal persister proportion, where $\partial V/\partial y = 0$. In Eq. S6 we approximate this optimal persister proportion using the numerical solution. To do this, we find the persister proportion, $y$ that minimises the absolute value of the finite difference approximation to $\partial V/\partial y = \partial \Psi/\partial y$, given by Eq. S19. In Eq. S6 we show this as a function of the time, $s$, and the growth rate, $\mu_t = m(z)$.

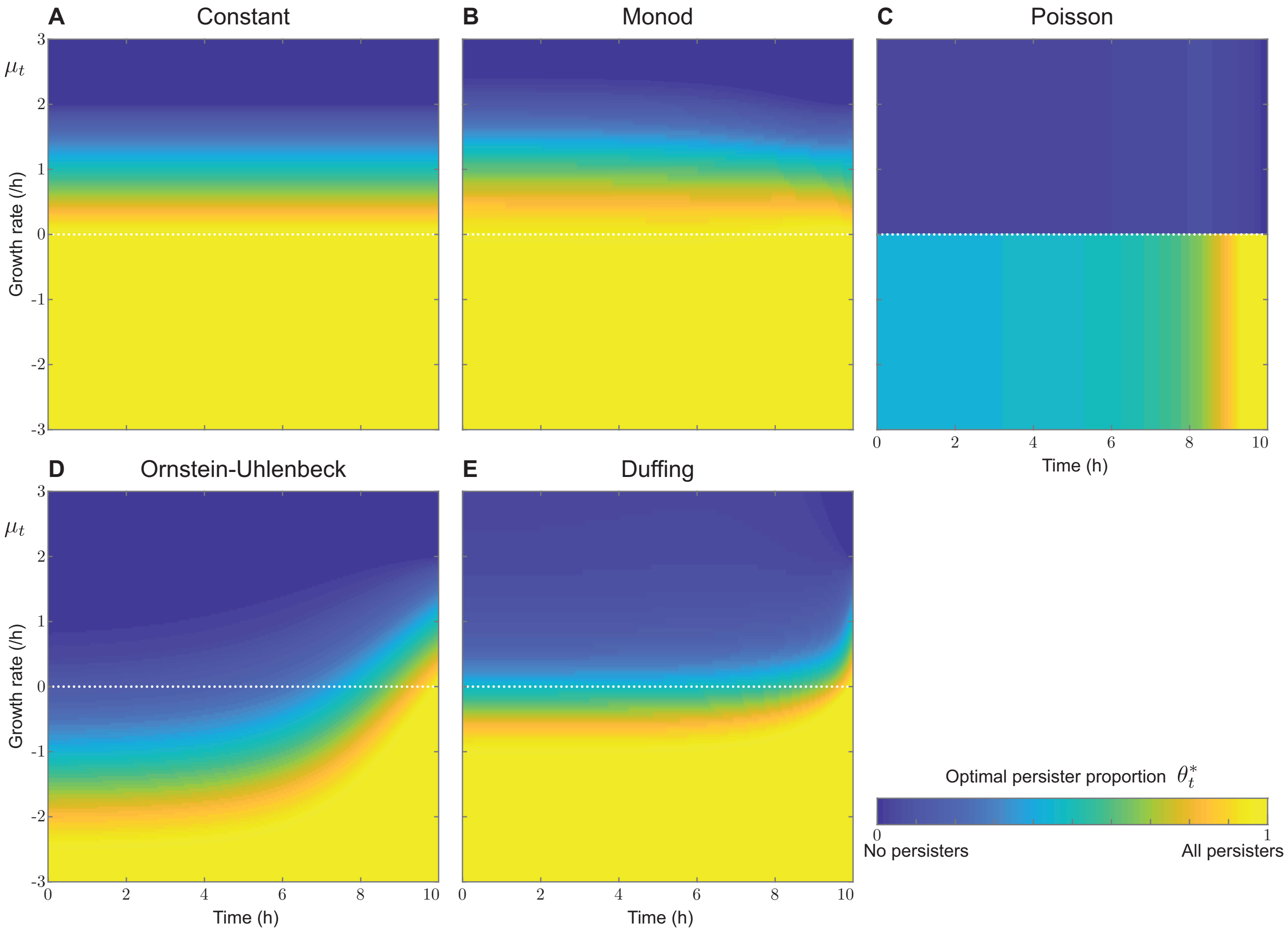

**Fig. S6.** Optimal persister proportion. The optimal persister proportion, as a function of time, $t$ (horizontal axis), and growth rate, $\mu_t$, for each environment. For (a), the optimal persister proportion is given by the analytical expression calculated in the main document. For (b)–(e), we consider the optimal persister proportion, $\theta_t^*$ to the value of $y$ that minimizes $\partial \Psi(y, z, t)/\partial y$, which we approximate using the numerical solution. In (c), the environment can only take two values $\zeta \in \{-1, 1\}$, which we assume corresponds to $\mu_t \geq 0$ and $\mu_t < 0$.
